# Pepsickle rapidly and accurately predicts proteasomal cleavage sites for improved neoantigen identification

**DOI:** 10.1101/2021.05.14.444244

**Authors:** Benjamin R. Weeder, Mary A. Wood, Ellysia Li, Abhinav Nellore, Reid F. Thompson

## Abstract

Proteasomal cleavage is a key component in protein turnover, as well as antigen presentation and subsequent immune response. Herein we present `pepsickle`, an open-source tool for proteasomal cleavage prediction with better *in vivo* prediction performance (AUC) and computational speed than current models available in the field, and with the ability to predict sites based on both constitutive and immunoproteasome profiles. Post-hoc filtering of predicted patient neoepitopes using `pepsickle` significantly enriches for immune-responsive epitopes and may represent a significant opportunity to improve current epitope prediction and vaccine development pipelines.

## Introduction

The constitutive proteasome is a multimeric protein complex that is best known for its role in the cleavage and recycling of cellular proteins marked for degradation ^1^. The proteasome also generates cleaved peptide fragments (epitopes) for immune surveillance via the major histocompatibility complex (MHC) class I antigen presentation pathway ^2^. This immune presentation functionality is critical for antiviral and other antimicrobial responses, and has particular relevance both in the setting of vaccine development and in a cancer context with the advent of immune checkpoint inhibitors ^3–7^.

Structurally, the proteasome consists of multiple subunits, a 20S barrel core housing the catalytic domains of the proteasome, and two 19S caps which aid in the unfolding of ubiquitin-tagged proteins ^1^. The barrel shape of the 20S core is derived from the fusion of four heptameric rings, the inner two of which contain a β1, β2, and β5 catalytic domain responsible for the cleavage of peptide bonds ^8^. Although all tissues express the constitutive proteasome, hematopoietic-lineage cells can also express the alternative catalytic domains β1i, β2i, and β5i in response to IFN-γ, which replace their analogues in the constitutive heptameric ring to form the immunoproteasome (Figure 1) ^9^. Previous studies support the presence of preferred cleavage motifs and differences in cleavage preferences between the immuno- and constitutive proteasomes; however, our understanding of how these preferences manifest is still not welldefined.^10,11^

**Figure 1.**
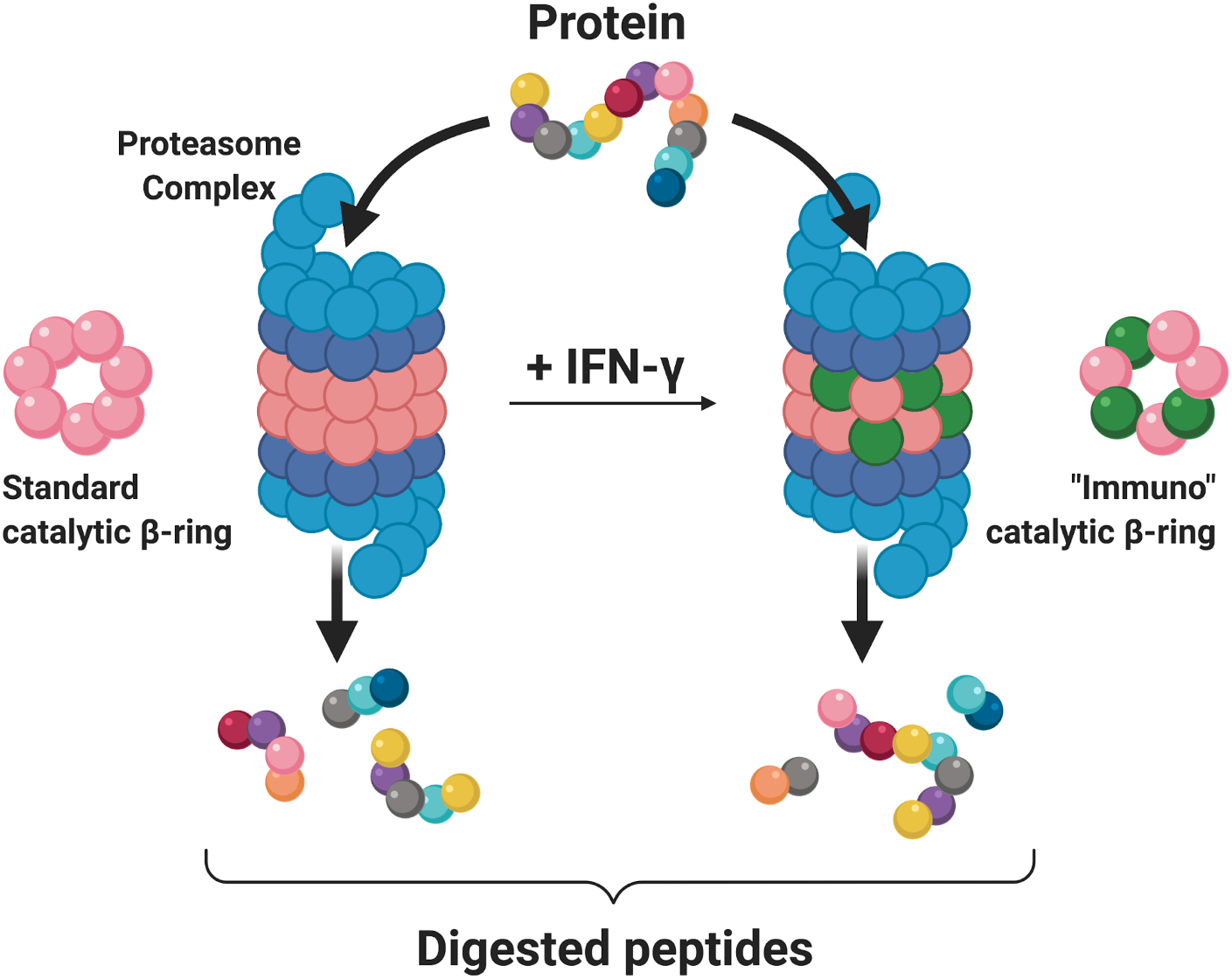
Protein degradation by the constitutive and immunoproteasome. Proteins trafficked to the proteasome complex are fed into the main 20S barrel, with the assistance of 19S caps (blue) that aid with unfolding and linearization. The catalytic domains of the standard ***β***-rings (pink) constituting the 20S barrel cleave the protein sequence and generate the resulting digested peptide fragments. In select tissues, exposure to interferon gamma (IFN-*γ*) results in replacement of the standard catalytic domains by alternative “immuno” catalytic domains (green). This transition in catalytic domain usage constitutes the construction of the immunoproteasome and may alter cleavage site preference. The differential digestion pattern of a single protein sequence (multi-colored) is depicted below the corresponding proteasome complex.

While existing tools to predict proteasomal cleavage sites are now widely adopted, each has significant limitations affecting the accuracy and/or scope of its predictions, with the potential for real world consequences (figure 2). NetChop 3.1, the most cited proteasomal cleavage tool, does not differentiate between constitutive and immuno-proteasomal cleavage when generating predictions ^10^. Further, tools such as the proteasomal cleavage prediction server (PCPS) provide options for predicting cleavage by the immunoproteasome but show poor model performance compared to NetChop when benchmarked ^10^. Finally, many available tools are either proprietary or otherwise unavailable to the public, complicating their use in both academic and industry analysis pipelines.

**Figure 2.**
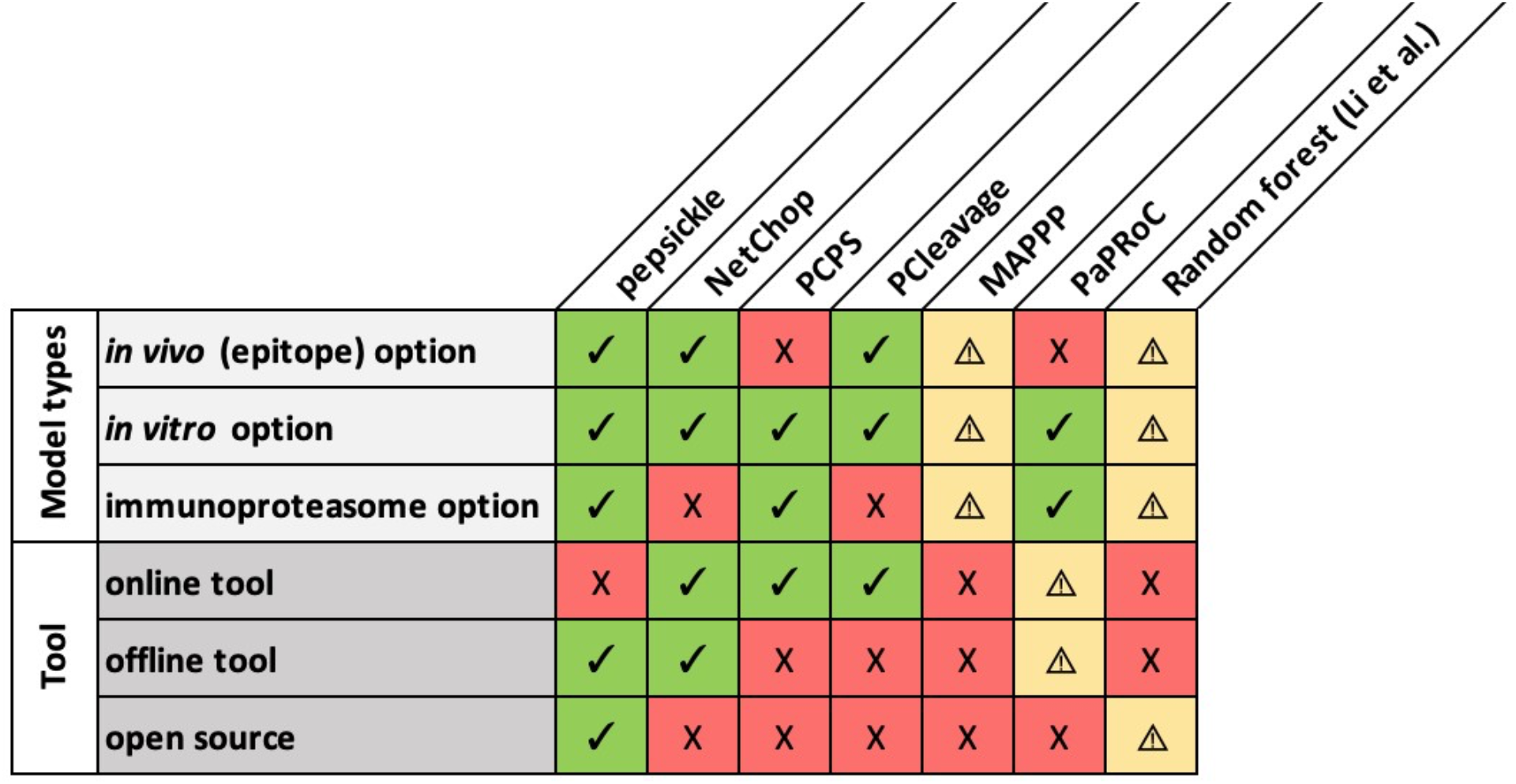
Comparison matrix of available proteasomal cleavage tools and their features. Eight proteasomal cleavage tools are shown (columns) along with their corresponding features (rows). Specific tools are as follows: `pepsickle` (presented here), NetChop 3.1 ^10^, the Proteasomal Cleavage Prediction Server (PCPS) ^64^, PCleavage ^65^, MAPPP ^66^, PAProC ^67^, and the random forest-based model described in Li et al. ^68^ Check marks (green) represent available features for each tool while X’s (red) represent unavailable features. Warning signs (yellow) represent missing information, or features that are mentioned but not currently available. For MAPPP, the referenced web server is no longer available and therefore we were unable to confirm tool features. For PaPRoC, we were unable to obtain the model despite repeated requests. For the random forest model proposed by Li et al., model weights for the proposed model are given, but source code is not available and the type of cleavage sites used (*in vivo vs. in vitro*) are undefined.

By leveraging a comprehensive set of proteasomal cleavage data and an ensemble based deep learning approach capable of representing complex motif preferences, we developed a set of models that consistently produce more accurate cleavage predictions than existing tools regardless of proteasomal context. We have deployed these models as an open-source command-line tool (`pepsickle`) for broad reuse and application.

## Methods

### Collection and processing of *in vitro* digestion map data for training and testing

We performed a literature search for all studies containing publicly available primary data from *in vitro* digestion experiments using 20S proteasomes. As the proteasome is highly conserved among mammalian species, digestion product results from non-human mammalian proteasomes were also included along with human-specific datasets ^11,12^. Search terms were “proteasome”, “proteasomal”, “cleavage”, “digestion”, “immunoproteasome”, “20S”, “i20S”, both alone and in various combinations. Ultimately, we identified 36 studies with relevant data (Table 1), from which we manually extracted individual cleavage sites, along with the parent peptide sequences from which they were derived^13,14,23–32,15,33–42,16,43–47,17–22^. Proteasome types present in the observed system (constitutive, immunoproteasome, or mixed) were also annotated for each cleavage experiment. Data from six 20S studies with unique source proteins were held out for downstream validation, while the remaining data (from 30 studies) were aggregated for model training and testing (Table 1). For *in vitro* digestion peptide fragments, both the N-terminal and C-terminal cleavage sites were used as cleavage examples. For each cleavage example, a context window was generated with the cleavage residue (C-terminus of the peptide fragment) as the “central” amino acid plus an equal number of upstream and downstream amino acids (Figure 8). Independent datasets with window sizes of 7 amino acids (3 upstream and 3 downstream from the central cleavage residue) and 21 amino acids (10 upstream and 10 downstream) were generated to allow for model optimization based on window size. Only unique cleavage windows were retained, yielding a total of 1,758 windows that are 7 amino acids in length and 1,819 windows that are 21 amino acids in length.

**Table 1.**
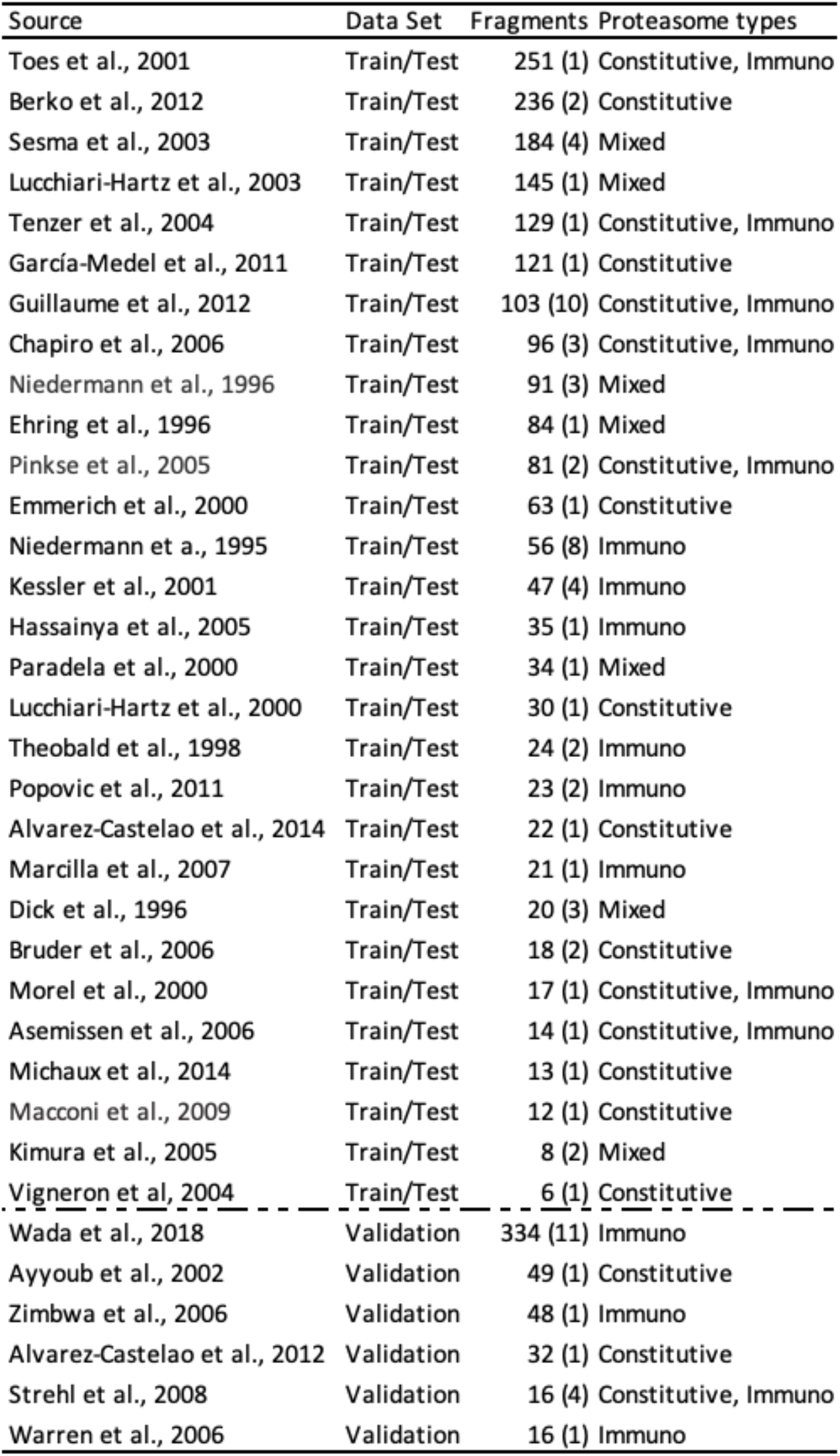
Summary of *in vitro* data. All data used for training, testing, and validating *in vitro* models is summarized above. Fragments represent the number of cleavage by-products reported in each primary literature source, with the number in parentheses representing the number of whole proteins or predigestion protein fragments used in each study. Proteasome type(s) denotes what proteasome was queried during experimentation with “constitutive” and/or “immuno” denoting isolated contexts, while “mixed” denotes testing in a non-isolated/heterogenous proteasomal context.

To generate companion non-cleavage examples for modeling, internal sites from each reported peptide fragment were considered as candidates. As above, 7 amino acid and 21 amino acid windows were generated for each non-cleavage example, and subsequently filtered to remove any duplicate sequences or overlaps with the set of non-cleavage examples. Before these candidates were included as null examples in the dataset, they were further filtered against all positive windows generated across all other studies, controlling for proteasome type, so that positive and negative cleavage examples were mutually exclusive.

### Collection and processing of epitope data for training and testing

To study *in vivo* cleavage sites, we extracted endogenously processed T-cell epitopes from three independent public databases (The Immune Epitope Database (IEDB) ^48^, AntiJen ^49^, and SYFPEITHI ^50^) as well as two primary literature sources: Bassani-Sternberg et al. (2016) and Rozanov et al. (2018). Data from Bassani-Sternberg et al. was maintained separately and used for downstream validation, while all other sources were aggregated for training and testing purposes. We restricted attention to mammalian naturally processed and presented peptide ligands of the MHC class I pathway. Epitopes were filtered to retain only those with an unambiguous substring position among known source protein sequence(s). Centered windows were generated around each C-terminal cleavage example as above, but using the full series of balanced window sizes from 7 amino acids to 21 amino acids, given the larger scale of the data and its accompanying power to detect significant differences in model performance. Only unique cleavage window sequences were retained, resulting in a total of 357,253 unique epitopes with C-terminal cleavage events. Note that epitope N-termini were not processed as cleavage examples due to the uncertainty resulting from N-terminal trimming by endoplasmic reticulum aminopeptidases (ERAP) ^53^.

To perform non-cleavage site inference, we first sampled internal amino acids for each epitope (Figure 9). Because the overwhelming majority of proteasomal digestion products have a length of at least two amino acids ^54–56^ and because peptides may be threaded into the proteasome in either the N- or Cterminal directions ^14^, we excluded 1 N-terminal and 1 C-terminal amino acid residue of each epitope from consideration in the potential non-cleavage data. As above, windows were generated for each remaining amino acid position for each potential window size. These potential non-cleavage windows were then filtered to remove any identical sequence matches within the set of positive cleavage examples from above. To additionally account for uncertainty in N-terminal cleavage position(s) due to ERAP ^53^, we removed any sequence matches with the set of windows generated by positions from the N-terminus of an epitope up to 16 amino acids upstream of each epitope’s C-terminus. Only unique non-cleavage window sequences were retained.

### Feature encoding

A vector of the following features was generated for each amino acid across windows in the cleavage and non-cleavage example sets (Supplementary Table S8). Amino acid identity was one-hot encoded as a bit vector of size 20, with each bit representing one of the standard amino acids. The null character (*) was used for padding, with all values as zero, while ambiguous amino acids were encoded as relevant combinations of non-zero values corresponding to their ambiguous components (e.g. B represents either aspartic acid or asparagine). Physical properties of amino acids were encoded as follows: side chain polarity was recorded as its isoelectric point (pI) ^57^, the molecular volume of each side chain was recorded as its partial molar volume at 37°C ^58^, the hydrophobicity of each side chain was characterized by its simulated contact angle with nanodroplets of water ^59^, and conformational entropy was derived from peptide bond angular observations among protein sequences without observed secondary structure (e.g. alpha helix) ^60^. Proteasomal context was also included where relevant (i.e., *in vitro* digestion data) as a single categorical feature with “C” representing the constitutive proteasome, “I” representing the immuno-proteasome, and “M” representing mixed systems with both proteasome types expressed.

### Gradient boosted decision tree structure and training

All gradient boosted classification models were implemented using the Scikit-learn package (v0.22.1) ^61^ for Python version 3.7. The aggregated positive and negative cleavage examples were randomly split to retain 80% of the examples for training and the remaining 20% for model testing. For each model, inversely balanced class weights were used, and the ‘RandomizedSearchCV’ class was used to determine the best option for the ‘max_features’ parameter (chosen from ‘auto’, ‘sqrt’, or ‘log2’) and the ‘n_estimators’ parameter (chosen from values of 100-1000, by 100) of the ‘GradientBoostingClassifier’ class. Randomized 10-fold cross validation was run for all combinations of parameters, and the best model (as determined by the ‘best_estimator_’ attribute) was retained. Model performance was evaluated based on AUC, using the pROC library (v1.16.2) ^62^ in R version 4.0.2.

Two distinct classification models were trained. One was based on the one hot encoded amino acid sequence identities, and another on the physical/chemical property encodings as previously described, normalized using the ‘fit_transform’ method of Scikit-learn’s ‘MinMaxScaler’ class. For epitope data, models were trained on amino acid window sizes 7 through 21 in length and compared to identify the model with optimal performance. For *in vitro* data, only the minimum (7) and maximum (21) length window libraries were assessed, accounting for constitutive v. immuno-proteasomal context.

### Neural network structure and training

All neural network models were implemented using the PyTorch package (version 1.3.1) ^63^ for Python version 3.7. The aggregated positive and negative cleavage examples were randomly split to retain 80% of the examples for training and the remaining 20% for model testing. We next trained two distinct cleavage classification models based on the proteasome type and either 1) amino acid identity encodings, or 2) amino acid physical property encodings as described previously. Each model consisted of an input layer, two hidden layers, and an output layer (Supplementary Figure S1). For all non-output layers, we applied batch normalization and a 20% dropout layer during each successive forward pass to improve model training and reduce overfitting. ReLU activation functions were employed at each step except for the output layer, where a softmax function was applied prior to final output. For the physical property-based model an additional convolutional layer (1D convolution with a 3 amino acid window and 1 amino acid step size) was applied to each physical property independently prior to passing values to the rest of the model. Cross entropy loss was used for backpropagation during training, with inverse class weights to account for class imbalance in the training set. Both models were trained for 36 epochs before training was halted, with AUC assessed on a new subset of the test data after each epoch and compared to the performance at the previous epoch, with the best performing model saved for downstream analysis.

For the two best-performing models (one identity-based and one based on physical properties), final testing performance was then assessed using a consensus approach, where the predicted probability of a test window representing a cleavage site was taken as the average probability across both models. For epitope data, models were trained on window sizes of 7 through 21 amino acids and subsequently compared to identify the window size with optimal performance. Due to the relatively small size of the *in vitro* dataset, only the minimum (7) and maximum (21) length window libraries were assessed, with additional information on constitutive vs. immuno-proteasomal context included as in the same manner used for the gradient boosted approach.

### Analysis of sampled feature space

To qualitatively assess how well our training data represented the broader space of possible peptides, we identified all unique 21 amino acid windows within the human proteome (https://www.uniprot.org/proteomes/UP000005640). Using these windows as background, we compared the shared UMAP space calculated with the first 10 principal components across the human proteome, as well as both *in vitro* and *in vivo* training sets using the four chemical properties at each amino acid position within the window, described previously, as the input feature set (Figure 6). Furthermore, we compared the sampling density for both datasets compared to the human background across the first 4 principal components to demonstrate the distribution of sampling in our training sets (Supplementary Figure S2). In addition to plots comparing the sampling space based on chemical properties, we also generated logo plots based on the amino acid frequencies for positive and negative examples in each training set (Supplementary Figure S5; Supplementary Figure S6). These plots were generated using the ultimate window sizes retained in modeling, 7 amino acids for *in vitro* data and 17 amino acids for *in vivo* data.

### Collection and processing of *in vitro* digestion map data for validation

Data from six 20S studies was held out from previous steps to be used in validation. In order to accommodate the analysis of performance for models using multiple different window sizes, window lengths of 21 amino acids were generated for each validation set cleavage example using reported peptide fragments and their source protein contexts. Companion non-cleavage windows were generated in the same way as before, with the exception that only one internal site was sampled at random during non-cleavage window generation to create an initially balanced set of positives and negatives. For validation windows, the additional step of filtering all validation windows with a non-unique interior window of 7 amino acids (smallest centered window size for the models to be assessed) was also taken to ensure no redundant windows were present in the validation set for any training window sizes from 7 to 21 amino acids. These windows were then screened against all training and testing examples to only retain unique, never before seen, entries in their respective sets. Ultimately, this generated 171 constitutive 20S cleavage windows and 54 immuno-proteasome 20S cleavage windows.

### Collection and processing of epitope data for validation

Data from Bassani-Sternberg et al. ^51^ was held out from previous steps to be used in validation. As described above, 21 amino acid windows were generated using reported epitopes and their source protein contexts. These windows were then screened to only retain unique entries in their respective sets and companion non-cleavage examples were generated as described previously, with the exception that only one internal site was sampled at random during non-cleavage window generation to create an initially balanced set of positives and negatives as with the in-vitro validation set. For validation windows, the additional step of filtering all validations windows with a non-unique interior window of 7 amino acids (smallest centered window size for the models to be assessed) was also taken to ensure no redundant windows were present in the validation set for any training window sizes from 7 to 21 amino acids. Finally, each window was also screened against those used in the training or testing sets to ensure none had been previously seen by the trained models. Ultimately, this generated 7,951 cleavage windows for validation.

### Model implementation and availability

Our *in vivo* and *in vitro* cleavage models were implemented in Python version 3.7. All deep learning models were generated using PyTorch version 1.3.1 ^63^, while all machine learning models were generated using Scikit-learn version 0.22.1 ^61^. The full instructions and code for replication of the analyses contained herein can be found at [https://github.com/pdxgx/pepsickle-paper], while the fully deployed command line version of ‘`pepsickle`’, along with relevant installation instructions can be found at [https://github.com/pdxgx/pepsickle]; `pepsickle` is open source and available under the MIT user license.

### Comparison of cleavage prediction tools

A literature search and browser query were performed to identify currently available tools for proteasomal cleavage prediction (search terms included “cleavage prediction”, “proteasomal prediction”, “cleavage prediction tool”, and “proteasomal cleavage prediction”). Through this search, six tools were identified including: NetChop 3.1 ^10^, the Proteasomal Cleavage Prediction Server (PCPS) ^64^, PCleavage ^65^, MAPPP ^66^, PAProC ^67^, and the random forest-based model described in Li et al. (2012). NetChop version 3.1 was downloaded (http://www.cbs.dtu.dk/services/NetChop/) and installed as a command line tool on a Linux server running CentOS 7.7.1908. Cleavage windows were given in FASTA format with predictions saved and assessed only for the point of potential cleavage in each window. PCPS was run via its web server implementation (http://imed.med.ucm.es/Tools/pcps/) with both constitutive proteasome and immunoproteasome options selected. For each *in vitro* data type, the model corresponding to the proteasome type was used, with only the midpoint of each window reported and recorded as described above. For *in vivo* epitope windows, both models were assessed in the same fashion, with results reported for the model achieving the best AUC. PCleavage was also run via its web server implementation (http://crdd.osdd.net/raghava/pcleavage/), however validation assessment was only performed for *in vivo* epitope windows using the default threshold of 0.3. For both constitutive proteasome and immunoproteasome data; PCleavage did not accept windows that spanned the C- or N- termini of a given source protein, which reduced the validation set size substantially and prevented paired comparisons with the other models available. However, three cleavage prediction tools were ultimately not functional in our hands: the MAPPP server is no longer available and we were unable to locate a publicly downloadable version of the tool; we were unable to obtain a working copy of PAProC II despite repeated requests; and we were unable to locate any web server or public tool implementing the model from Li et al..

Computational performance was assessed for all tools not reliant on a web server (i.e., NetChop 3.1, `pepsickle`). Using a dedicated node (Intel Xeon E5-2697 v2 2.70GHz, single thread mode) on a Linux server running CentOS 7.7.1908, both *in vitro* and epitope-based models for NetChop 3.1 and `pepsickle` were applied to a performance test set consisting of all proteins in the human proteome (https://www.uniprot.org/proteomes/UP000005640). Total CPU times were calculated as the user ‘time’ + ‘sys’ time for each prediction model (Supplementary Table S7).

### Model cross-comparison assessments

The cross performance of both constitutive-based and immuno-based *in vitro* models were assessed using the same *in vivo* validation set used for our epitope trained model (Supplementary Table S9). Cleavage predictions were generated using the internal 7 amino acid window centered within each larger 21 amino acid validation window. Predictions were reported at the same central amino acid with the default prediction probability threshold of 0.5 used for determining cleavage vs. non-cleavage predictions. Reasoning that positive epitope cleavage examples should be predicted with good accuracy, while negative examples would not due to the complex selectivity of the downstream antigen processing and presentation pathway (e.g., MHC binding), only the percentage of correctly captured positive cleavage examples was assessed (sensitivity). This removes the possibility of misreporting true cleavage events that are filtered during post-cleavage processing as model misclassifications.

### Collection of patient-derived immune response data and model application

Three primary literature articles including patient specific predicted tumor neoepitopes and epitope-specific immune responses were identified for model application, including: 1) the Ott et al. patient-specific melanoma vaccine study ^69^, 2) the MuPeXI neoepitope prediction study ^70^, and 3) a large scale neoepitope prediction comparison from the Tumor Neoantigen Selection Alliance (TESLA) ^71^. From sources 1 and 2 where gene/protein sources for each predicted epitope were provided, each mutated candidate was mapped back to its original proteomic position to retrieve upstream (10 amino acids) and downstream (10 amino acids) contexts. For predicted neoepitopes within 10 amino acids from the start or end of the protein, positions were buffered using “*” prior to model input. For predicted neoepitopes reported in the TESLA study, original source proteins were not provided; instead, candidate neoepitope sequences were queried against the human reference proteome using the BLAST ^72^ command line tool with the following parameters: -matrix BLOSUM62, -evalue 200000, -comp_based_stats F. Only ungapped alignments were retained, allowing for a singular mismatch at the mutated position with exact matches at all other positions. Protein contexts around each candidate neoepitope were generated as described for the other two studies, however all candidate neoepitopes resulting in more than one unique context window were filtered out to remove any candidate neoepitopes with an ambiguous source in the proteome. All predicted neoepitopes across the three studies were also annotated as “responsive” or “non-responsive” based on the reported patient specific immune response. This resulted in 762 candidate neoepitopes, of which 45 (5.9%) were reported as inducing a patient-specific immune response.

After all context windows were collected, the *in vivo* `pepsickle` model was applied to the Cterminal position of each proposed neoepitope candidate returning the predicted C-terminal cleavage probability. Median cleavage probabilities for predicted neoepitopes that elicited a patient specific immune response were compared to those that were predicted but for which an immune response was not verified using a Wilcoxon ranked sum test. Additionally, the use of cleavage probability as a classification threshold was assessed using the 25th percentile of predicted cleavage probabilities across all candidate neoepitopes as a cutoff. The proportion of responsive vs. non-responsive neoepitopes that were properly identified using this thresholding approach was assessed using a Chi-square(df) test for independence.

## Results

### *In vitro* digestion-based cleavage prediction

We identified 36 publicly available *in vitro* digestion datasets, constituting both 20S constitutive and 20S immuno-proteasomal cleavage experiments (Table 1). From these, six studies were reserved for external validation (validation set), while the rest were aggregated to generate a training and testing dataset containing cleavage information from 1,984 peptide fragments generated across constitutive, immuno-, and mixed proteasomal contexts. We then trained a gradient boosted classifier based on windows of 7 amino acids in length centering on each cleaved site. Residues within the window were encoded as the physical properties (polarity, molecular volume, hydrophobicity, and conformational entropy) of each amino acid at each given position in the window. Using annotated proteasome types, the model was trained to differentiate between sites cleaved by the immuno-proteasome and those cleaved by the constitutive proteasome, returning the probability of cleavage at the center of each window. This model achieved a test set AUC of 0.759 (Supplementary Table S1). We explored whether additional peptide context around each cleavage site (21 amino acid windows) improved model performance; however, a comparison of models trained with both window sizes showed no increase in AUC when applied to testing the data (DeLong’s T-test, p = 0.558). Similarly, we assessed whether a more complex deep learning model could improve cleavage predictions. This also did not appear to increase performance significantly (p = 0.558). We therefore report and discuss the results from our 7 amino acid gradient boosted classifier (`pepsickle`) hereafter.

We next assessed *in vitro* `pepsickle` performance on an independent validation set, consisting of 171 constitutive proteasome and 54 immunoproteasome examples, respectively. Our model achieved an AUC of 0.821 on the constitutive proteasome validation set and 0.789 on the immuno-proteasome validation set, respectively. Using the same validation sets, we assessed the corresponding performance of existing tools including NetChop 3.1 and the Proteasomal Cleavage Prediction Server (PCPS) (figure 3). Note that PCleavage was omitted from these *in vitro* based comparisons due to its inability to process cleavage sites whose context windows span a peptide fragment’s N- or C-termini (54.4% of the constitutive and 81.5% of the immuno validation data respectively). We found that `pepsickle` has significantly higher predictive performance on constitutive proteasomal data compared to PCPS, but similar performance compared to NetChop 3.1 (Figure 3). When applied to immuno-proteasomal data, our model compared similarly to both PCPS and NetChop 3.1, acknowledging limited statistical power to detect a difference given the small sample size (Figure 4; Supplementary Table S2).

**Figure 3.**
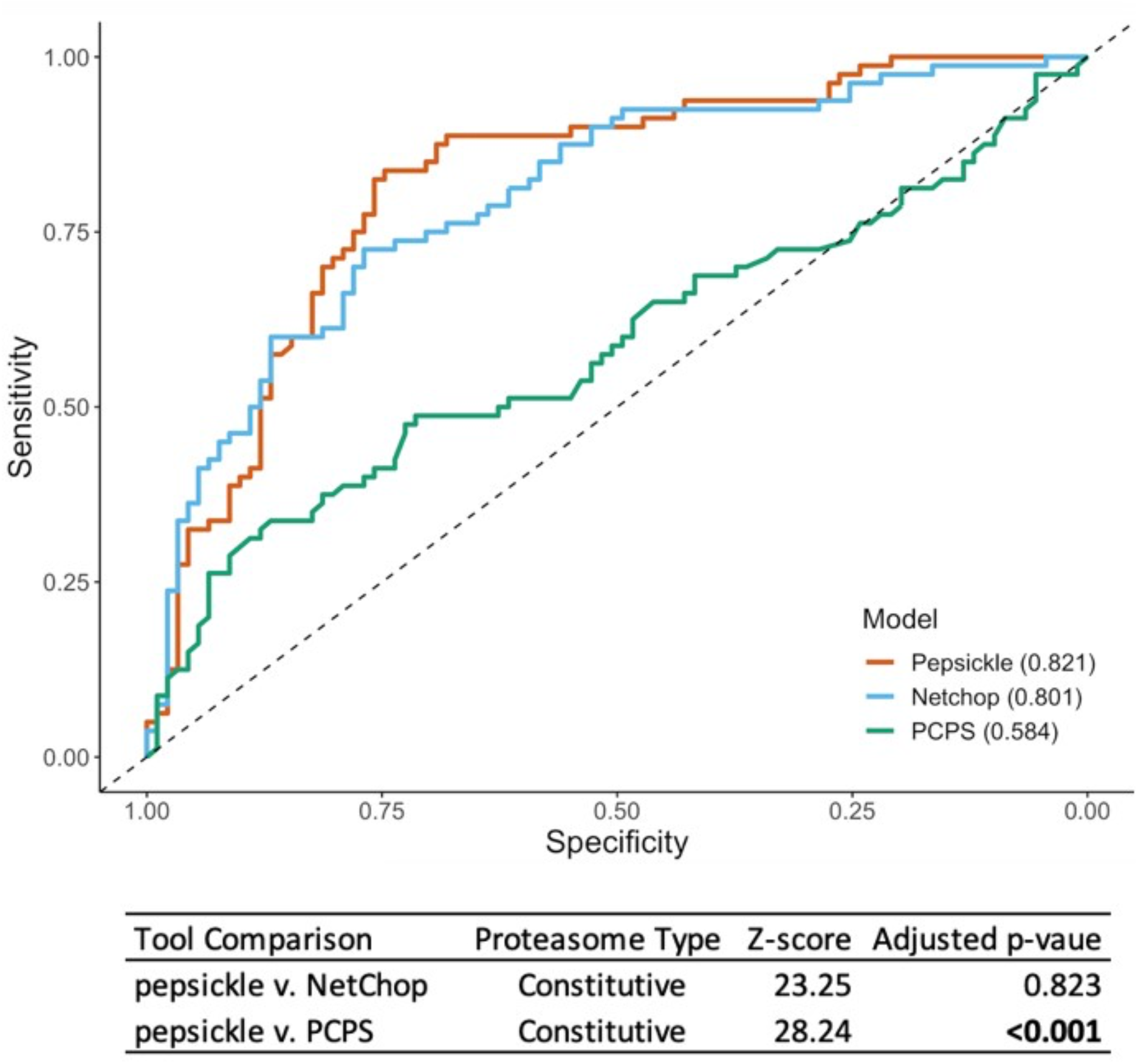
Performance comparison of cleavage prediction models on constitutive proteasome data. Receiver operating characteristic (ROC) curves are shown for each of three cleavage prediction models, as denoted in legend, with corresponding area under the curve (AUC) values reported in parentheses. Sensitivity (y-axis) and specificity (x-axis) were both evaluated using a validation set (n = 171) consisting of 80 cleavage and 91 non-cleavage *in vitro* examples not seen during the training or testing of our models (see Methods). For `pepsickle` (our model) and PCPS, the constitutive proteasome models with default settings were used. For NetChop 3.1, the *in vitro* model was used with default settings (no specification is available for proteasome type). PCleavage was omitted from this comparison due to restrictions on window sizes and the inability to process the full set of validation examples. Statistical pairwise comparisons of ROC curves (Delong’s tests) are shown in corresponding table values (Z-score), with significance reported as p-values after Benjamini-Hochberg correction for multiple comparisons.

**Figure 4.**
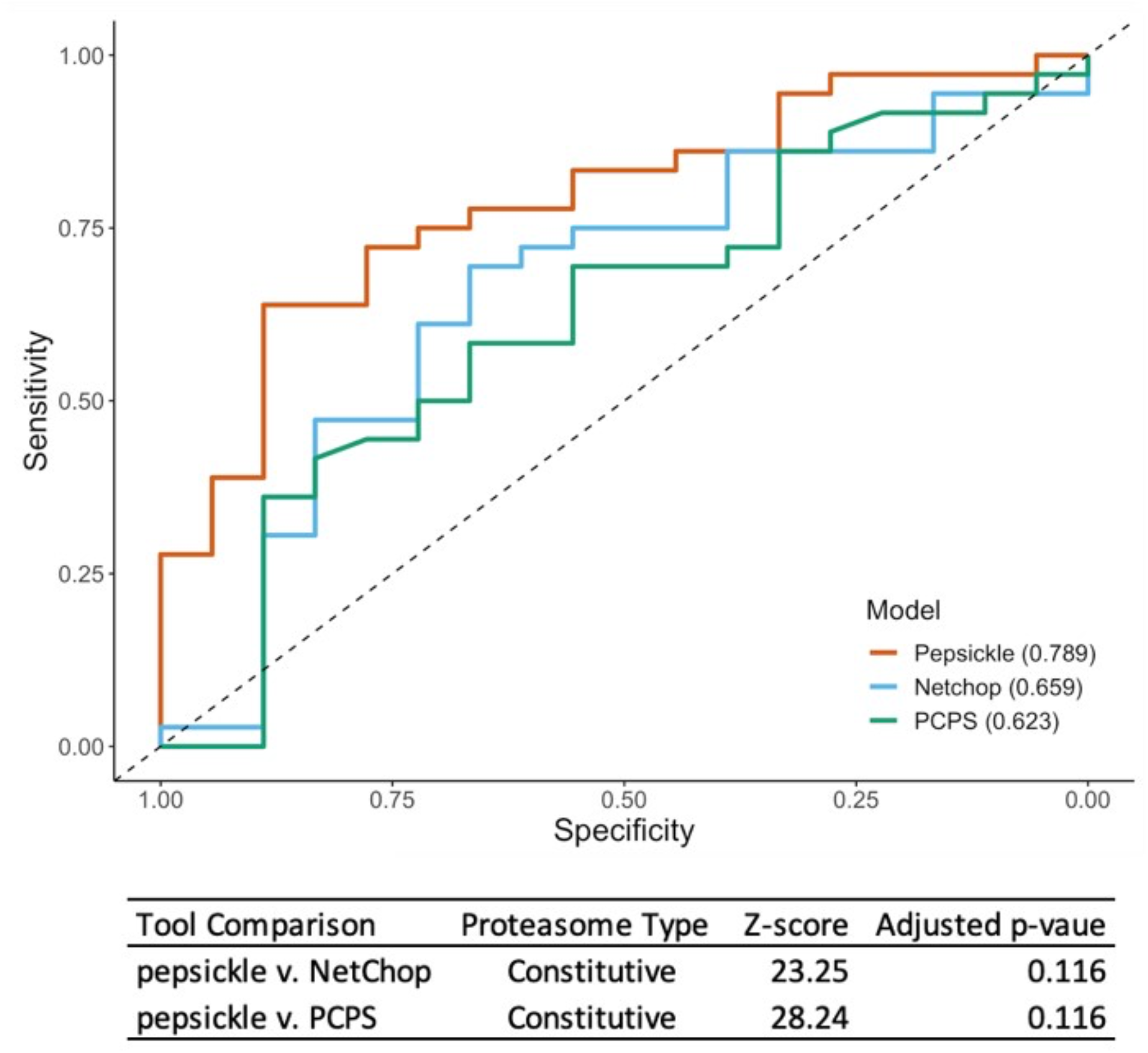
Performance comparison of cleavage prediction models on immunoproteasome data. Receiver operating characteristic (ROC) curves are shown for each of three cleavage prediction models, as denoted in legend, with corresponding area under the curve (AUC) values reported in parentheses. Sensitivity (yaxis) and specificity (x-axis) were both evaluated using a validation set (n = 54) consisting of 36 cleavage and 18 non-cleavage in-vitro examples not seen during the training or testing of our models (see Methods). For `pepsickle` (our model) and PCPS, the immunoproteasome models with default settings were used. For NetChop 3.1, the *in vitro* model was used with default settings (no specification is available for proteasome type). PCleavage was omitted from this comparison due to restrictions on window sizes and the inability to process the full set of validation examples. Statistical pairwise comparisons of ROC curves (Delong’s tests) are shown in corresponding table values (Z-score), with significance reported as pvalues after Benjamini-Hochberg correction for multiple comparisons.

### Epitope-based cleavage prediction

To better interrogate *in vivo* proteasomal cleavage, we identified 357,253 naturally processed human and mammalian class I epitopes from publicly available data (Table 2). Using a deep learning framework, we trained a consensus based neural network on amino acid sequence and physical properties to predict epitope C-terminal cleavage events, independent of proteasome type (Supplementary Figure S1). Performance of our deep learning model was compared across all odd window sizes ranging from 7 amino acids to 21 amino acids, with windows centered on the cleavage site as described above (Supplementary Table S4). When applied to testing data, the model trained on 17 amino acid windows performed significantly better than the model trained on 7 amino acid windows; however increasing window size beyond 17 amino acids did not improve performance further. We additionally studied the influence of model complexity, finding that the consensus-based deep learning approach performed better than a more simplistic random forest model trained using the same window size (DeLong’s T-test, p = 0.021). We therefore report and discuss results from the consensus-based model (`pepsickle`) using 17 amino acid windows hereafter.

**Table 2.** Summary of Epitope Data Sources. Sources of epitope data used for training, testing, and validation.

| Source | Data Set | Fragments Reported |
| --- | --- | --- |
| Immune Epitope Database (IEDB)<sup>54</sup> | Train/Test | 498,419 |
| SYFPEITHI Database<sup>56</sup> | Train/Test | 4,433 |
| Rozanov et al.<sup>58</sup> | Train/Test | 3,254 |
| Antijen Database<sup>55</sup> | Train/Test | 1,492 |
| Bassani-Sternberg et al.<sup>57</sup> | Validation | 99,356 |

We assessed performance of the `pepsickle` epitope model on an independent validation set. This dataset consisted of 7,951 examples not replicated in either the training or testing datasets used previously. When applied to this validation data, our deep learning-based ensemble net achieved an AUC of 0.878, representing a significant improvement in AUC over the corresponding performances of existing tools including NetChop 3.1, PCPS, and PCleavage collectively (Figure 5). In addition, `pepsickle` showed better recall and F-1 score than the other models compared (Supplementary Table S5).

**Figure 5.**
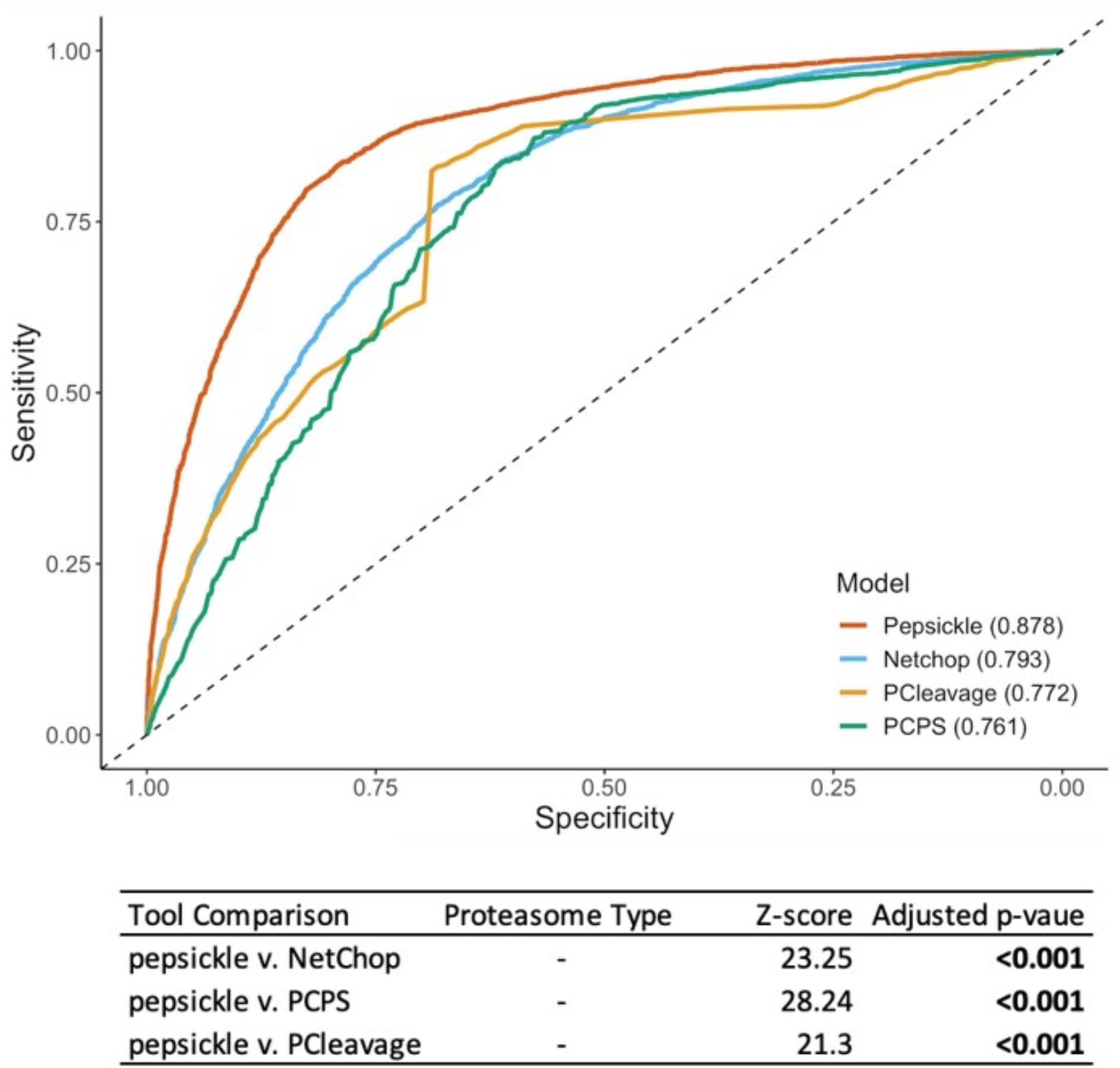
Performance comparison of cleavage prediction models on epitope data. Receiver operating characteristic (ROC) curves are shown for each of four cleavage prediction models, as denoted in legend, with corresponding area under the curve (AUC) values reported in parentheses. Sensitivity (y-axis) and specificity (x-axis) were both evaluated using a validation set (n = 7,951) consisting of 3,566 cleavage and 4,385 non-cleavage epitope examples not seen during the training or testing of our models (see Methods). Default epitope-based models were used for `pepsickle` (our model), Netchop 3.1, and PCleavage predictions, while constitutive model predictions from the default model 1 were used for PCPS (PCPS immuno-proteasome predictions were inferior and therefore omitted). Statistical pairwise comparisons of ROC curves (Delong’s tests) are shown in corresponding table values (Z-score), with significance reported as p-values after Benjamini-Hochberg correction for multiple comparisons.

### Computational performance of pepsickle

In addition to predictive ability, we also assessed the computational speed of `pepsickle`, for both *in vitro* and epitope-based cleavage predictions. Using a list of all protein sequences in the human proteome as a benchmark dataset (n=113,576, including all isoforms and computationally predicted sequences), `pepsickle` was able to achieve a total processing time of 154m 46s for *in vitro* predictions (approximately 124 milliseconds per 1,000 predictions) and 158m 21s for epitope predictions (approximately 127 milliseconds per 1,000 predictions) (Supplementary Table S7). These times were compared to NetChop 3.1, run in an identical, controlled computing environment. We found that `pepsickle` is 68.5% faster for *in vitro* cleavage predictions (154m 46s v. 260m 50s) and 242% faster for epitope-based predictions (158m 21s v. 542m 40s) compared to NetChop 3.1.

### *In vitro* digestion and *in vivo* epitope-based models differ in prediction performance, but with similar feature importance

Given the substantial differences between sources of training data for both *in vitro* digestion and *in vivo* epitope-based models, we next sought to evaluate commonalities in the learned feature sets by evaluating cross-performance of our *in vitro* model on epitope validation data. Acknowledging that epitope-based data is implicitly subject to multiple components of the antigen processing pathway following proteasomal cleavage^73^, we evaluated the accuracy of the *in vitro* model exclusively on positive cleavage examples from the *in vivo* epitope validation set (i.e., all positive examples must have necessarily undergone proteasomal cleavage). Based on this metric, our *in vitro* constitutive model was able to correctly identify 69.9% of the cleavage events observed in the epitope validation set, while our immunoproteasome model was able to correctly identify 54.5%. Performance by both *in vitro* models on this data is substantially lower than the performance of the original epitope-based model, which was able to capture 82.8% of true cleavage events.

Because cross-data assessments for both *in vitro* models represent a substantial performance decrease compared to assessment on like-kind data, we sought to further qualify the distinct commonalities and differences between *in vitro* digestion and *in vivo* epitope-based datasets. Using 21 amino acid windows, we compared both training sets to a set of all possible 21 amino acid windows from the human proteome. By overlapping UMAP projections of the windows sampled in the *in vivo* epitope set with those generated by the human proteome, we were able to visually demonstrate that the majority of the sample space constituted by the human proteome was sampled; however substantial portions were not sufficiently sampled in the *in vitro* dataset (FIGURE 6). Similarly, the underlying density distribution for samples in the *in vitro* dataset differed substantially from that seen in both the *in vivo* dataset and human proteome background (Supplementary Figure S2).

**Figure 6.**
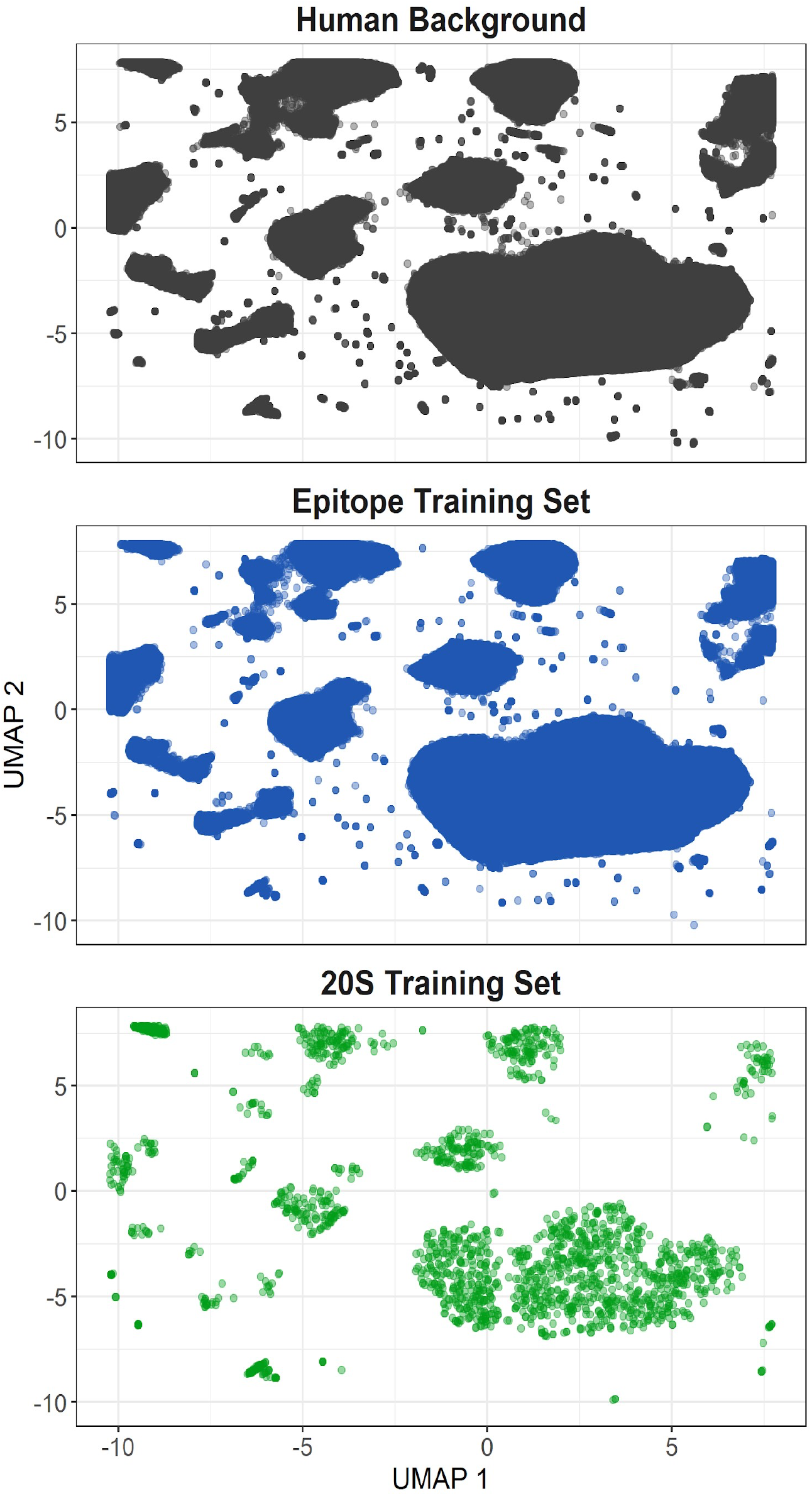
Training set projections in UMAP space. Amino acid windows (21 residues long) were generated for the whole human proteome (gray), all epitope training examples (blue), and all 20S training examples (green). Principle components were generated from the physical properties of each amino acid at each window position. UMAP projections were generated from the first 10 principal components.

To further investigate whether differences in the training set representations altered the learned features for each prediction model, we plotted the feature importance for both our *in vivo* and *in vitro* models (Supplementary Figure S3; Supplementary Figure S4). Acknowledging that model weights are not directly comparable, we found that similar patterns of amino acid physical properties identified cleavage sites across both models: in particular, molecular volume and hydrophobicity at the C-terminal amino acid, as well as the relatively higher importance of conformational entropy at the ‘1 position and polarity at the ‘2 position compared to other features at the same locations.

### Proteasomal cleavage predicts epitope-specific immune responses -

We next assessed the potential additive contribution of our model to predicting epitope-specific immune responses in real-world patient data. We identified 762 candidate epitopes from three studies with extensive immunoprofiling data: 1) the Ott et al. patient-specific melanoma vaccine study ^69^, 2) the MuPeXI neoepitope prediction study ^70^, and 3) a large scale neoepitope prediction benchmarking effort from the Tumor Neoantigen Selection Alliance (TESLA) ^71^. From these studies, we identified 45 epitopes that elicited an immune response, as well as 717 non-responsive epitopes.

Using the `pepsickle` epitope-based cleavage model, we predicted C-terminal cleavage probability for all predicted epitopes regardless of corresponding immune response status (FIGURE 7). We demonstrated that the median terminal cleavage probability is significantly higher for immune responsive epitopes compared to those that were predicted but did not elicit an immune response (Wilcoxon ranked sum test, p = 0.036). Despite the heavy pre-selection of these epitopes using a collection of predictive methodologies, we find that `pepsickle` based cleavage prediction (≥ 25th percentile threshold) significantly enriched the proportion of immune responsive epitope candidates with 40% of responsive vs. 24.4% of non-responsive candidates falling in the top quartile (*χ*_1_^2^= 4.86, p = 0.027). This represents a 59.6% increase in the positive predictive value after cleavage-based filtering. Notably, we find that cleavage predictions for two studies follow the trend seen in the aggregate data ^69,71^, but the third study does not ^70^. While this heterogeneity warrants further investigations, these findings suggest that even when used as a post-hoc filter, `pepsickle` based cleavage predictions may help improve the identification of patient-specific, immune-responsive neoepitopes.

**Figure 7.**
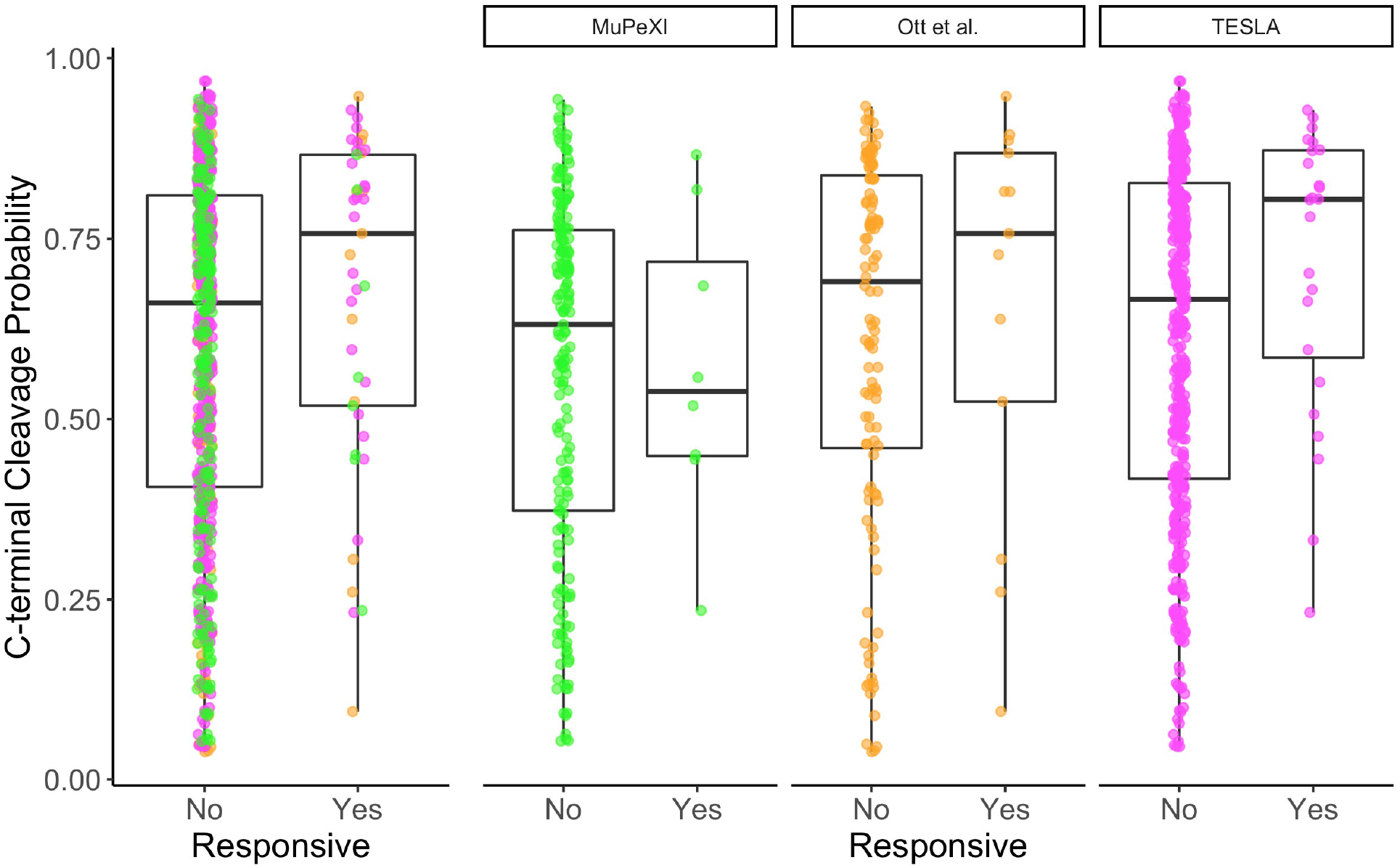
C-terminal cleavage predictions on patient neoepitopes. Predicted neoepitopes from three studies were accumulated. Cleavage predictions at the C-terminus of each epitope were generated using `pepsickle` and plotted, with predicted epitopes divided based on whether or not they elicited an immune response. Box plot results are shown in aggregate (left) as well as on a per-study basis (right) with values indicating median (horizontal black lines), 25-75%ile (box), and range (“whiskers”), and with colors corresponding to the study origin (green = MuPeXI, orange = Ott et al., magenta = TESLA).

**Figure 8.**
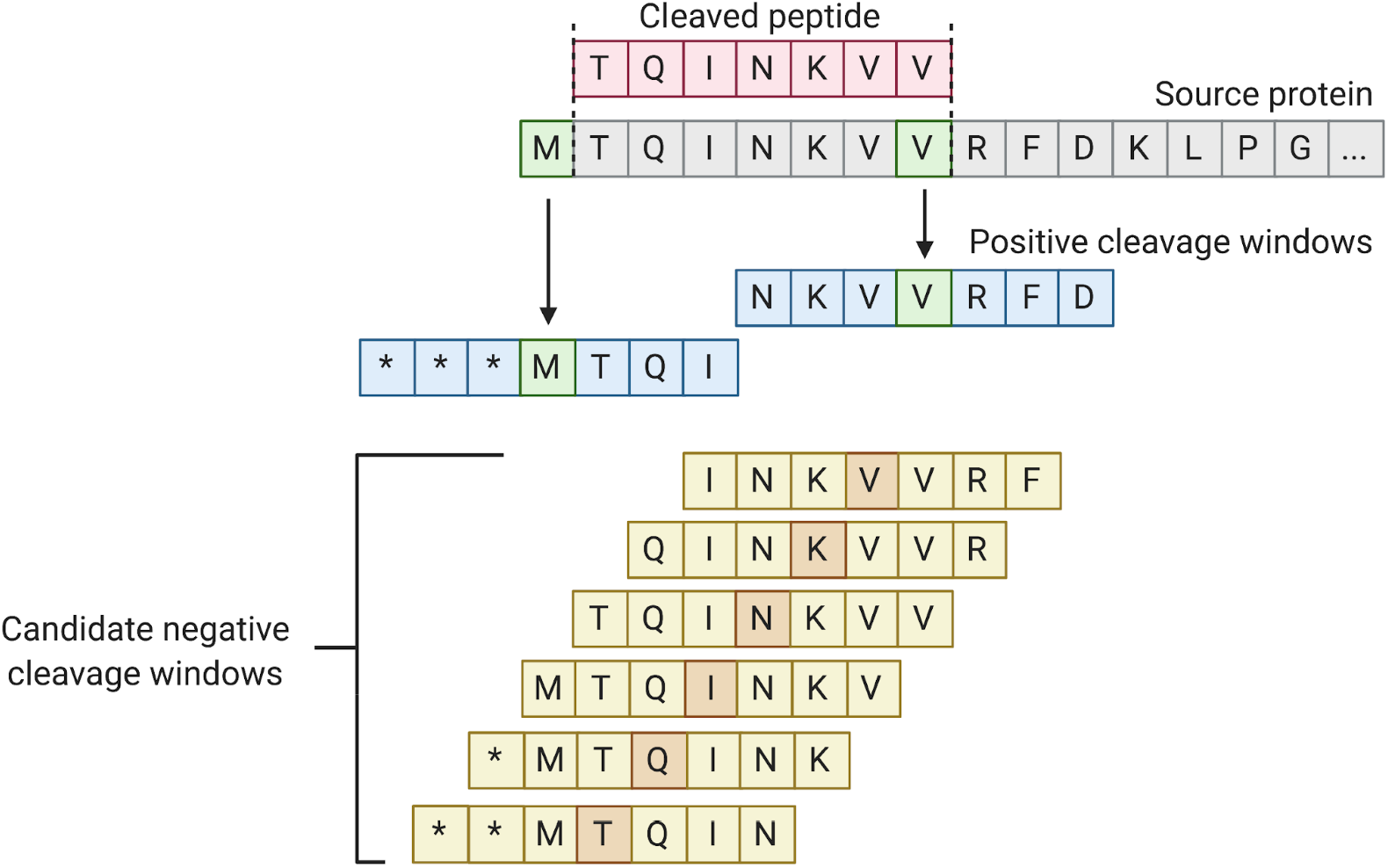
Generation of the *in vitro* dataset. Each identified cleaved peptide fragment (red) was mapped back to its source sequence (gray). Using the C-terminus of the fragment, as well as the amino acid prior to the N-terminus of the fragment as cleavage sites (green, with each of their respective downstream bonds cleaved by the proteasome), cleavage windows (blue) were generated using 3 amino acids upstream and downstream of the cleavage sites identified. Candidate non-cleavage windows (yellow) were generated using the same windowed approach on internal amino acids within the epitope. Before candidate negatives were included in the dataset, they were screened against all positive identified cleavage sites from both N- and C-termini of reported fragments. Note that * indicates the lack of an amino acid (i.e. amino acid position is beyond the peptide terminus).

**Figure 9.**
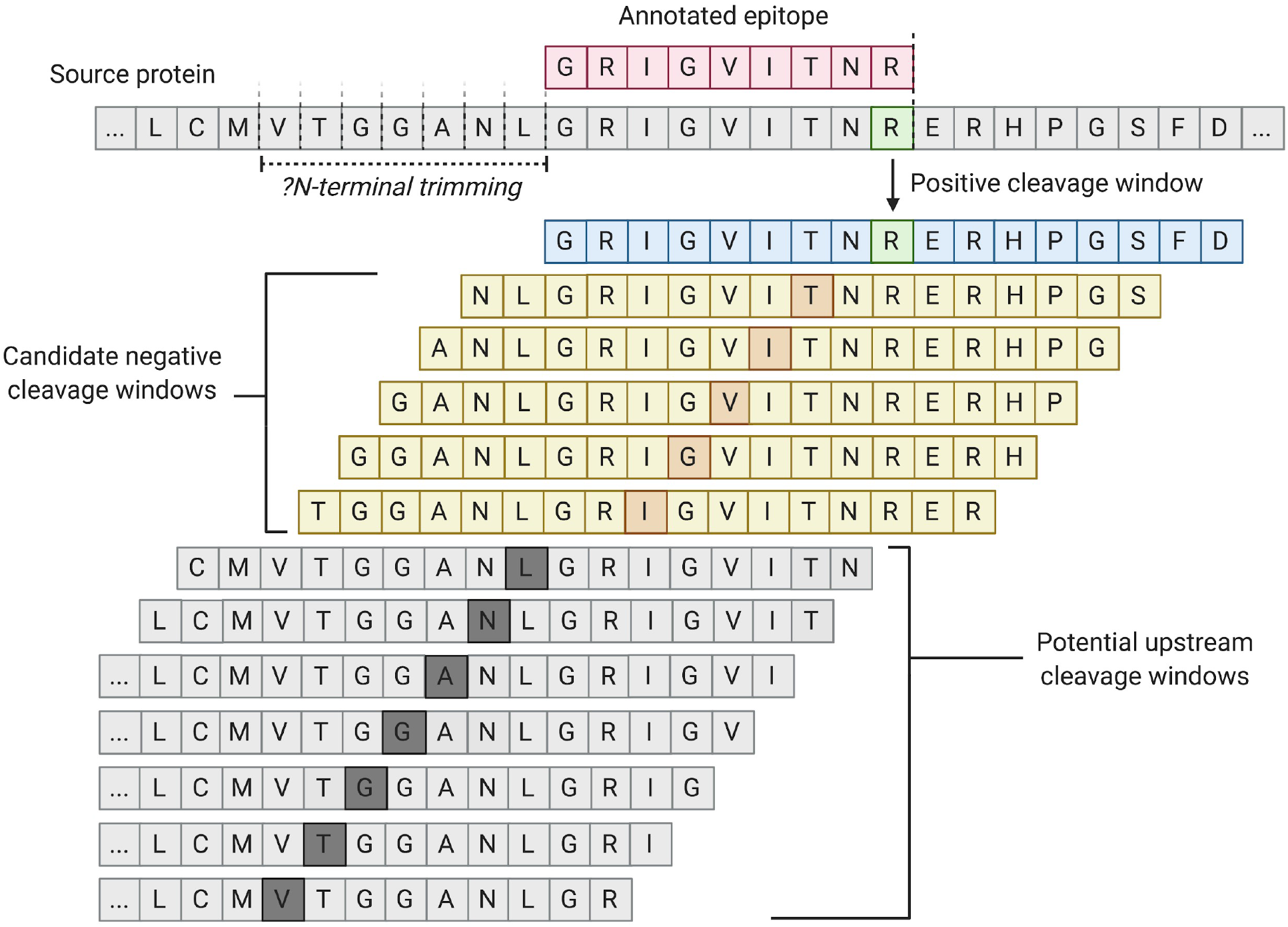
Generation of the epitope dataset. Each identified epitope (red) was mapped back to its source sequence. Using the C-terminus of the epitope as the cleavage site (green), with the downstream bond as the one cleaved by the proteasome), cleavage windows (blue) were generated using 8 amino acids up and downstream from the site identified. Candidate non-cleavage windows (yellow) were generated using the same windowed approach on internal amino acids within the epitope, with the exclusion of the first two and last two amino acids which served as a buffer region to account for minimum proteasomal fragment size. Before candidate negatives were included in the dataset, they were screened against all positive identified cleavage sites as well as against a set of potential upstream cleavage sites (gray); generated by using the same windowed approach on the upstream window that could encapsulate the N-terminal cleavage site prior to ERAP trimming.

## Discussion

To the best of our knowledge, the data aggregated for this study represents the largest compilation of *in vitro* and *in vivo* cleavage events to date. Applying machine and deep learning techniques to this data, we have improved upon the current state of the field by developing an *in vivo* model of proteasomal cleavage prediction with improved performance (AUC) over currently available tools. In addition, we have created an *in vitro* model with performance comparable to the current best-in-class model, NetChop 3.1, but with significantly decreased computational costs and the ability to differentiate between immunoproteasome and standard proteasome cleavage profiles. Although further investigation is needed, application of our *in vivo* model to patient-derived neoepitope data suggests that including cleavage information in epitope prediction may improve novel target identification and may be a key component missing from the majority of current prediction tools. This is consistent with recent evidence demonstrating the value of incorporating proteasomal cleavage predictions into epitope prediction pipelines ^74^.

Despite `pepsickle`’s promising performance using both *in vivo* and *in vitro* models, we note several limitations to our work. The primary challenge given the structure of the *in vivo* data, is that non-cleavage events must be determined heuristically. Although we use stringent filtering criteria throughout our pipeline, accurate negative examples are reliant on sufficient sampling of true cleavage events and may be biased by lack of reporting for less studied portions of the proteome. While we did not see evidence that our models learned features unrelated to cleavage, such as MHC binding ^75^, it remains possible that these and other latent biological features may have been partially learned by our models in addition to true cleavage-specific features (Supplementary Figure S3; Supplementary Figure S4). Additionally, our *in vitro* models are based on relatively small datasets with heterogeneous experimental methodologies, and only a small subset cleanly evaluate the respective roles of the constitutive and immunoproteasomes on the same source proteins. For both *in vivo* and *in vitro* data, poor sampling from some regions of the proteome is also of concern, due at least in part to a scientific focus on proteins relevant for cancer and autoimmunity, as well as the experimental limitations of mass spectrometry ^76^. Finally, while the data suggests our model should perform well on previously unseen data, the discrepancies seen in model application to per-study immune response data raise questions of broader generalizability in certain applied contexts.

Pepsickle provides a promising, open-source, tool for proteasomal cleavage prediction, which may be implemented on its own or otherwise integrated into existing epitope prediction pipelines. Given the recent successes and increasing emphasis on developing and deploying mRNA-based vaccines for individual patients ^77^ and whole populations ^3,78^, any concrete improvements in the accuracy of these epitope prediction pipelines could carry transformative clinical value. We also note that an improved capacity to predict proteasomal cleavage could contribute to our understanding of protein turnover and recycling in healthy and diseased contexts ^79,80^ and lead to improvements in rational protein design ^81^.

The performance and potential of `pepsickle` described in this text are encouraging, however many questions remain unanswered. The heterogeneity of C-terminal cleavage profiles seen in our study specific `pepsickle` application raises the question of whether cleavage prediction is universally helpful in target identification, or if specific study or design contexts are required to see benefit from cleavage predictions. In addition, whether or not there is an impact of using proteasomal prediction ad-hoc (via integration with existing neoepitope prediction pipelines ^82,83^) vs. post-hoc remains to be seen. Application of `pepsickle` to more patient derived data in the future will help us better understand the broader potential of applying cleavage prediction in this space, with the potential for broad implications in the research and clinical communities.

## Data Access

Source code is available at https://github.com/pdxgx/pepsickle-paper under the Massachusetts Institute of Technology (MIT) license, including data extracted from primary literature (https://github.com/pdxgx/pepsickle-paper/tree/master/data/raw) and scripts for parsing public databases mentioned herein (https://github.com/pdxgx/pepsickle-paper/tree/master/scripts/database_pulls).

## Competing Interest Statement

The authors of this manuscript report no competing interests.

## Notes

https://github.com/pdxgx/pepsickle

https://github.com/pdxgx/pepsickle-paper

### Competing Interest Statement

The authors have declared no competing interest.

## References

1. Ciechanover, A. The ubiquitin-proteasome proteolytic pathway. Cell 79, 13–21 (1994).

2. Kloetzel, P. M. & Ossendorp, F. Proteasome and peptidase function in MHC-class-I-mediated antigen presentation. Curr. Opin. Immunol. 16, 76–81 (2004).

3. Pardi, N., Hogan, M. J., Porter, F. W. & Weissman, D. mRNA vaccines-a new era in vaccinology. Nat. Rev. Drug Discov. 17, 261–279 (2018).

4. Sijts, E. J. A. M. & Kloetzel, P. M. The role of the proteasome in the generation of MHC class I ligands and immune responses. Cell. Mol. Life Sci. 68, 1491–1502 (2011).

5. Ribas, A. & Wolchok, J. D. Cancer immunotherapy using checkpoint blockade. Science (80-.). 359, 1350–1355 (2018).

6. Carretero, R. et al. Regression of melanoma metastases after immunotherapy is associated with activation of antigen presentation and interferon-mediated rejection genes. Int. J. Cancer 131, 387–395 (2012).

7. Ostrand-Rosenberg, S. Tumor immunotherapy: the tumor cell as an antigen-presenting cell. Curr. Opin. Immunol. 6, 722–727 (1994).

8. Adams, J. The proteasome: Structure, function, and role in the cell. Cancer Treat. Rev. 29, 3–9 (2003).

9. Budenholzer, L., Cheng, C. L., Li, Y. & Hochstrasser, M. Proteasome Structure and Assembly. J. Mol. Biol. 429, 3500–3524 (2017).

10. Nielsen, M., Lundegaard, C., Lund, O. & Keşmir, C. The role of the proteasome in generating cytotoxic T-cell epitopes: Insights obtained from improved predictions of proteasomal cleavage. Immunogenetics 57, 33–41 (2005).

11. Fort, P., Kajava, A. V., Delsuc, F. & Coux, O. Evolution of proteasome regulators in Eukaryotes. Genome Biol. Evol. 7, 1363–1379 (2015).

12. Tomko, R. J. & Hochstrasser, M. Molecular architecture and assembly of the eukaryotic proteasome. Annu. Rev. Biochem. 82, 415–445 (2013).

13. Toes, R. E. M. et al. Discrete cleavage motifs of constitutive and immunoproteasomes revealed by quantitative analysis of cleavage products. J. Exp. Med. 194, 1–12 (2001).

14. Berko, D. et al. The Direction of Protein Entry into the Proteasome Determines the Variety of Products and Depends on the Force Needed to Unfold Its Two Termini. Mol. Cell 48, 601–611 (2012).

15. Sesma, L., Alvarez, I., Marcilla, M., Paradela, A. & López De Castro, J. A. Species-specific Differences in Proteasomal Processing and Tapasin-mediated Loading Influence Peptide Presentation by HLA-B27 in Murine Cells. J. Biol. Chem. 278, 46461–46472 (2003).

16. Lucchiari-Hartz, M. et al. Differential proteasomal processing of hydrophobic and hydrophilic protein regions: Contribution to cytotoxic T lymphocyte epitope clustering in HIV-1-Nef. Proc. Natl. Acad. Sci. U. S. A. 100, 7755–7760 (2003).

17. Tenzer, S. et al. Quantitative Analysis of Prion-Protein Degradation by Constitutive and Immuno-20S Proteasomes Indicates Differences Correlated with Disease Susceptibility. J. Immunol. 172, 1083–1091 (2004).

18. García-Medel, N., Sanz-Bravo, A., Barnea, E., Admon, A. & López De Castro, J. A. The origin of proteasome-inhibitor resistant HLA class I peptidomes: A Study with HLA-A*68:01. Mol. Cell. Proteomics 11, (2012).

19. Guillaume, B. et al. Analysis of the Processing of Seven Human Tumor Antigens by Intermediate Proteasomes. J. Immunol. 189, 3538–3547 (2012).

20. Chapiro, J. et al. Destructive Cleavage of Antigenic Peptides Either by the Immunoproteasome or by the Standard Proteasome Results in Differential Antigen Presentation. J. Immunol. 176, 1053–1061 (2006).

21. Niedermann, G. et al. The proteolytic fragments generated by vertebrate proteasomes: Structural relationships to major histocompatibility complex class I binding peptides. Proc. Natl. Acad. Sci. U. S. A. 93, 8572–8577 (1996).

22. Ehring, B., Meyer, T. H., Eckerskorn, C., Lottspeich, F. & Tampé, R. Effects of majorhistocompatibility-complex-encoded subunits on the peptidase and proteolytic activities of human 20S proteasomes: Cleavage of proteins and antigenic peptides. Eur. J. Biochem. 235, 404–415 (1996).

23. Pinkse, G. G. M. et al. Autoreactive CD8 T cells associated with β cell destruction in type 1 diabetes. Proc. Natl. Acad. Sci. U. S. A. 102, 18425–18430 (2005).

24. Emmerich, N. P. N. et al. The human 26 S and 20 S proteasomes generate overlapping but different sets of peptide fragments from a model protein substrate. J. Biol. Chem. 275, 21140–21148 (2000).

25. Niedermann, G. et al. Contribution of proteasome-mediated proteolysis to the hierarchy of epitopes presented by major histocompatibility complex class I molecules. Immunity 2, 289–299 (1995).

26. Kessler, J. H. et al. Efficient identification of novel HLA-A*0201-presented cytotoxic T lymphocyte epitopes in the widely expressed tumor antigen PRAME by proteasome-mediated digestion analysis. J. Exp. Med. 193, 73–88 (2001).

27. Hassainya, Y. et al. Identification of naturally processed HLA-A2 - Restricted proinsulin epitopes by reversed immunology. Diabetes 54, 2053–2059 (2005).

28. Paradela, A. et al. Limited Diversity of Peptides Related to an Alloreactive T Cell Epitope in the HLA-B27-Bound Peptide Repertoire Results from Restrictions at Multiple Steps Along the Processing-Loading Pathway. J. Immunol. 164, 329–337 (2000).

29. Lucchiari-Hartz, M. et al. Cytotoxic T lymphocyte epitopes of HIV-1 Nef: Generation of multiple definitive major histocompatibility complex class I ligands by proteasomes. J. Exp. Med. 191, 239–252 (2000).

30. Theobald, M. et al. The sequence alteration associated with a mutational hotspot in p53 protects cells from lysis by cytotoxic T lymphocytes specific for a flanking peptide epitope. J. Exp. Med. 188, 1017–1028 (1998).

31. Popović, J. et al. The only proposed T-cell epitope derived from the TEL-AML1 translocation is not naturally processed. Blood 118, 946–954 (2011).

32. Alvarez-Castelao, B., Goethals, M., Vandekerckhove, J. & Castaño, J. G. Mechanism of cleavage of alpha-synuclein by the 20S proteasome and modulation of its degradation by the RedOx state of the N-terminal methionines. Biochim. Biophys. Acta - Mol. Cell Res. 1843, 352–365 (2014).

33. Marcilla, M., De Castro, J. A. L., Castaño, J. G. & Alvarez, I. Infection with Salmonella typhimurium has no effect on the composition and cleavage specificity of the 20S proteasome in human lymphoid cells. Immunology 122, 131–139 (2007).

34. Dick, T. P. et al. Coordinated dual cleavages induced by the proteasome regulator PA28 lead to dominant MHC ligands. Cell 86, 253–262 (1996).

35. Bruder, D. et al. Multiple synergizing factors contribute to the strength of the CD8+ T cell response against listeriolysin O. Int. Immunol. 18, 89–100 (2006).

36. Morel, S. et al. Processing of some antigens by the standard proteasome but not by the immunoproteasome results in poor presentation by dendritic cells. Immunity 12, 107–117 (2000).

37. Asemissen, A. M. et al. Identification of a highly immunogenic HLA-A*01-binding T cell epitope of WT1. Clin. Cancer Res. 12, 7476–7482 (2006).

38. Michaux, A. et al. A Spliced Antigenic Peptide Comprising a Single Spliced Amino Acid Is Produced in the Proteasome by Reverse Splicing of a Longer Peptide Fragment followed by Trimming. J. Immunol. 192, 1962–1971 (2014).

39. Macconi, D. et al. Proteasomal processing of albumin by renal dendritic cells generates antigenic peptides. J. Am. Soc. Nephrol. 20, 123–130 (2009).

40. Kimura, Y., Gushima, T., Rawale, S., Kaumaya, P. & Walker, C. M. Escape Mutations Alter Proteasome Processing of Major Histocompatibility Complex Class I-Restricted Epitopes in Persistent Hepatitis C Virus Infection. J. Virol. 79, 4870–4876 (2005).

41. Vigneron, N. et al. An Antigenic Peptide Produced by Peptide Splicing in the Proteasome. Science (80-.). 304, 587–590 (2004).

42. Wada, H. et al. Development of a novel immunoproteasome digestion assay for synthetic long peptide vaccine design. PLoS One 13, e0199249 (2018).

43. Ayyoub, M. et al. Proteasome-Assisted Identification of a SSX-2-Derived Epitope Recognized by Tumor-Reactive CTL Infiltrating Metastatic Melanoma. J. Immunol. 168, 1717–1722 (2002).

44. Zimbwa, P. et al. Precise Identification of a Human Immunodeficiency Virus Type 1 Antigen Processing Mutant. J. Virol. 81, 2031–2038 (2007).

45. Alvarez-Castelao, B. et al. Reduced protein stability of human DJ-1/PARK7 L166P, linked to autosomal recessive Parkinson disease, is due to direct endoproteolytic cleavage by the proteasome. Biochim. Biophys. Acta - Mol. Cell Res. 1823, 524–533 (2012).

46. Strehl, B. et al. Antitopes define preferential proteasomal cleavage site usage. J. Biol. Chem. 283, 17891–17897 (2008).

47. Warren, E. H. et al. An antigen produced by splicing of noncontiguous peptides in the reverse order. Science (80-.). 313, 1444–1447 (2006).

48. Vita, R. et al. The Immune Epitope Database (IEDB): 2018 update. Nucleic Acids Res. 47, D339–D343 (2019).

49. McSparron, H., Blythe, M. J., Zygouri, C., Doytchinova, I. A. & Flower, D. R. JenPep: A novel computational information resource for immunobiology and vaccinology. J. Chem. Inf. Comput. Sci. 43, 1276–1287 (2003).

50. Rammensee, H. G., Bachmann, J., Emmerich, N. P. N., Bachor, O. A. & Stevanović, S. SYFPEITHI: Database for MHC ligands and peptide motifs. Immunogenetics 50, 213–219 (1999).

51. Bassani-Sternberg, M. et al. Direct identification of clinically relevant neoepitopes presented on native human melanoma tissue by mass spectrometry. Nat. Commun. 7, 13404 (2016).

52. Rozanov, D. V. et al. MHC class I loaded ligands from breast cancer cell lines: A potential HLA-I-typed antigen collection. J. Proteomics 176, 13–23 (2018).

53. Evnouchidou, I. & van Endert, P. Peptide trimming by endoplasmic reticulum aminopeptidases: Role of MHC class I binding and ERAP dimerization. Hum. Immunol. 80, 290–295 (2019).

54. Köhler, A. et al. The axial channel of the proteasome core particle is gated by the Rpt2 ATPase and controls both substrate entry and product release. Mol. Cell 7, 1143–1152 (2001).

55. Kisselev, A. F., Akopian, T. N., Woo, K. M. & Goldberg, A. L. The sizes of peptides generated from protein by mammalian 26 and 20 S proteasomes. Implications for understanding the degradative mechanism and antigen presentation. J. Biol. Chem. 274, 3363–3371 (1999).

56. Nussbaum, A. K. et al. Cleavage motifs of the yeast 20S proteasome β subunits deduced from digests of enolase. Proc. Natl. Acad. Sci. U. S. A. 95, 12504–12509 (1998).

57. P, D. CRC Handbook of Chemistry and Physics. Journal of Molecular Structure 268, (CRC press, 1992).

58. Häckel, M., Hinz, H. J. & Hedwig, G. R. Partial molar volumes of proteins: Amino acid side-chain contributions derived from the partial molar volumes of some tripeptides over the temperature range 10-90°C. Biophys. Chem. 82, 35–50 (1999).

59. Zhu, C. et al. Characterizing hydrophobicity of amino acid side chains in a protein environment via measuring contact angle of a water nanodroplet on planarpeptide network. Proc. Natl. Acad. Sci. U. S. A. 113, 12946–12951 (2016).

60. Fogolari, F. et al. Distance-based configurational entropy of proteins from molecular dynamics simulations. PLoS One 10, e0132356 (2015).

61. Pedregosa, F. et al. Scikit-learn: Machine Learning in Python. J. Mach. Learn. Res. 12, 2825–2830 (2011).

62. Robin, X. et al. pROC: An open-source package for R and S+ to analyze and compare ROC curves. BMC Bioinformatics 12, 77 (2011).

63. Paszke, A. et al. PyTorch: An imperative style, high-performance deep learning library. arXiv 32, 12 (2019).

64. Gomez-Perosanz, M., Ras-Carmona, A. & Reche, P. A. PCPS: A web server to predict proteasomal cleavage sites. in Methods in Molecular Biology (ed. Tomar, N.) 2131, 399–406 (Springer US, 2020).

65. Bhasin, M. & Raghava, G. P. S. Pcleavage: An SVM based method for prediction of constitutive proteasome and immunoproteasome cleavage sites in antigenic sequences. Nucleic Acids Res. 33, W202–207 (2005).

66. Holzhütter, H. G. & Kloetzel, P. M. A kinetic model of vertebrate 20S proteasome accounting for the generation of major proteolytic fragments from oligomeric peptide substrates. Biophys. J. 79, 1196–1205 (2000).

67. Nussbaum, A. K., Kuttler, C., Hadeler, K. P., Rammensee, H. G. & Schild, H. PAProC: A prediction algorithm for proteasomal cleavages available on the WWW. Immunogenetics 53, 87–94 (2001).

68. Li, B. Q., Cai, Y. D., Feng, K. Y. & Zhao, G. J. Prediction of Protein Cleavage Site with Feature Selection by Random Forest. PLoS One 7, e45854 (2012).

69. Ott, P. A. et al. An immunogenic personal neoantigen vaccine for patients with melanoma. Nature 547, 217–221 (2017).

70. Bjerregaard, A. M., Nielsen, M., Hadrup, S. R., Szallasi, Z. & Eklund, A. C. MuPeXI: prediction of neoepitopes from tumor sequencing data. Cancer Immunol. Immunother. 66, 1123–1130 (2017).

71. Wells, D. K. et al. Key Parameters of Tumor Epitope Immunogenicity Revealed Through a Consortium Approach Improve Neoantigen Prediction. Cell 183, 818–834.e13 (2020).

72. Altschul, S. F., Gish, W., Miller, W., Myers, E. W. & Lipman, D. J. Basic local alignment search tool. J. Mol. Biol. 215, 403–410 (1990).

73. Rock, K. L., Reits, E. & Neefjes, J. Present Yourself! By MHC Class I and MHC Class II Molecules. Trends Immunol. 37, 724–737 (2016).

74. Sarkizova, S. et al. A large peptidome dataset improves HLA class I epitope prediction across most of the human population. Nat. Biotechnol. 38, 199–209 (2020).

75. Wieczorek, M. et al. Major histocompatibility complex (MHC) class I and MHC class II proteins: Conformational plasticity in antigen presentation. Front. Immunol. 8, (2017).

76. Danilova, Y., Voronkova, A., Sulimov, P. & Kertész-Farkas, A. Bias in False Discovery Rate Estimation in Mass-Spectrometry-Based Peptide Identification. J. Proteome Res. 18, 2354–2358 (2019).

77. Türeci, Ö. et al. Targeting the heterogeneity of cancer with individualized neoepitope vaccines. Clin. Cancer Res. 22, 1885–1896 (2016).

78. Maruggi, G., Zhang, C., Li, J., Ulmer, J. B. & Yu, D. mRNA as a Transformative Technology for Vaccine Development to Control Infectious Diseases. Mol. Ther. 27, 757–772 (2019).

79. Smail, M. A., Reigle, J. K. & McCullumsmith, R. E. Using protein turnover to expand the applications of transcriptomics. Sci. Rep. 11, 4403 (2021).

80. Bibo-Verdugo, B., Jiang, Z., Caffrey, C. R. & O’Donoghue, A. J. Targeting proteasomes in infectious organisms to combat disease. FEBS J. 284, 1503–1517 (2017).

81. Coluzza, I. Computational protein design: A review. J. Phys. Condens. Matter 29, 143001 (2017).

82. Wood, M. A., Weeder, B. R., David, J. K., Nellore, A. & Thompson, R. F. Burden of tumor mutations, neoepitopes, and other variants are dubious predictors of cancer immunotherapy response and overall survival. bioRxiv 12, 33 (2020).

83. Hundal, J. et al. PVACtools: A computational toolkit to identify and visualize cancer neoantigens. Cancer Immunol. Res. 8, 409–420 (2020).

