## Supplementary_tables for "Pepsickle rapidly and accurately predicts proteasomal cleavage sites for improved neoantigen identification"

**Table S1. In vitro model performances by window size.**

| Model 1 | | | | Model 2 | | | | |
|---|---|---|---|---|---|---|---|---|
| feature input | Size | AUC | | feature input | Size | AUC | delta-AUC | adj-pvalue |
| **chemical** | **7** | **0.759** | **vs.** | **sequence** | **7** | **0.723** | **0.036** | **0.002** |
| **chemical** | **21** | **0.771** | **vs.** | **sequence** | **21** | **0.743** | **0.028** | **0.012** |
| chemical | 7 | 0.759 | vs. | chemical | 21 | 0.771 | -0.012 | 0.558 |
| sequence | 7 | 0.723 | vs. | sequence | 21 | 0.743 | -0.020 | 0.513 |

Comparison of model performances on in vitro test data based on feature window size used for trianing input.

**Table S2. Epitope sequence-based deep learning layer sizes by input window size.**

| | Layers | | | | |
|---|---|---|---|---|---|
| Window size | Input | internal #1 | internal #2 | internal #3 | Output |
| 7 | 140 | 136 | 68 | 34 | 2 |
| 9 | 180 | 136 | 68 | 34 | 2 |
| 11 | 220 | 136 | 68 | 34 | 2 |
| 13 | 260 | 136 | 68 | 34 | 2 |
| 15 | 300 | 136 | 68 | 34 | 2 |
| 17 | 340 | 136 | 68 | 34 | 2 |
| 19 | 380 | 136 | 68 | 34 | 2 |
| 21 | 420 | 136 | 68 | 34 | 2 |

Model layer sizes for sequence based deep learning models based on training window size. For all internal layers a ReLU activation function was used with 20% dropout. For the final output layer, log(Softmax) was used as the activation function.

**Table S3. Epitope chemical-based deep learning layer sizes by input window size.**

| | Layers | | | | |
|---|---|---|---|---|---|
| Window size | input layer | lutional layer | internal #1 | internal #2 | Output |
| 7 | 28 | 20 | 38 | 20 | 2 |
| 9 | 36 | 28 | 38 | 20 | 2 |
| 11 | 44 | 36 | 38 | 20 | 2 |
| 13 | 52 | 44 | 38 | 20 | 2 |
| 15 | 60 | 52 | 38 | 20 | 2 |
| 17 | 68 | 60 | 38 | 20 | 2 |
| 19 | 76 | 68 | 38 | 20 | 2 |
| 21 | 84 | 76 | 38 | 20 | 2 |

Model layer sizes for physical property based deep learning models based on training window size. For the initial convolutional layer, a 1D convolution with span=3 and step=1 was used. For all internal layers a ReLU activation function was used with 20% dropout and for the final output layer, log(Softmax) was used as the activation function.

**Table S4. Immunoproteasome validation set power analysis.**

| | Estimated Delta-AUC | # cases | # controls | N | Beta | Alpha |
|---|---|---|---|---|---|---|
| Current Beta, controlled alpha | 0.130 | 36 | 18 | 54 | 0.665 | 0.05 |
| Target Beta, controlled alpha | 0.130 | 55 | 27 | 82 | 0.800 | 0.05 |

Statistical power analysis for `pepsickle` and NetChop 3.1 *in vitro* model comparison on immunoproteasome validation data. Initial estimate represents the actual type II error based on the available *in vitro* immunoproteasome data with type I error controlled at 0.05 and the observed difference in AUC. The second estimate represents the requisite number of cases and controls required to achieve a target type II error (1 - β) of 0.20, using the same case/control ratio and AUC difference observed in the available validation set.

**Table S5. Epitope model test-set comparisons by window size.**

| Model Window Size | | Model Comparison (large - small) | |
|---|---|---|---|
| Small | Large | AUC difference | Adj. P-value |
| 7 | 9 | -0.010 | 0.352 |
| 7 | 11 | -0.003 | 0.778 |
| 7 | 13 | 0.002 | 0.851 |
| 7 | 15 | 0.017 | 0.080 |
| **7** | **17** | **0.022** | **0.019** |
| **7** | **19** | **0.030** | **0.001** |
| **7** | **21** | **0.029** | **0.001** |
| 9 | 11 | 0.007 | 0.527 |
| 9 | 13 | 0.012 | 0.269 |
| **9** | **15** | **0.027** | **0.004** |
| **9** | **17** | **0.031** | **0.001** |
| **9** | **19** | **0.040** | **<0.001** |
| **9** | **21** | **0.038** | **<0.001** |
| 11 | 13 | 0.005 | 0.655 |
| **11** | **15** | **0.020** | **0.034** |
| **11** | **17** | **0.025** | **0.007** |
| **11** | **19** | **0.033** | **<0.001** |
| **11** | **21** | **0.032** | **0.001** |
| 13 | 15 | 0.015 | 0.111 |
| **13** | **17** | **0.020** | **0.028** |
| **13** | **19** | **0.028** | **0.001** |
| **13** | **21** | **0.027** | **0.002** |
| 15 | 17 | 0.005 | 0.624 |
| 15 | 19 | 0.013 | 0.136 |
| 15 | 21 | 0.012 | 0.189 |
| 17 | 19 | 0.008 | 0.352 |
| 17 | 21 | 0.007 | 0.450 |
| 19 | 21 | -0.001 | 0.851 |

Epitope models were trained on odd size starting windows between 7 amino acids and 21 amino acids. Each model was applied to the held out test set and assessed based on AUC. Delong's tests were used for pairwise statistical comparisons of the performance for each size of base window in contrast with other window sizes. P-values for comparisons were adjusted using Benjamini-Hochberg p-value correction and model comparisons with significant differences in AUC after correction are denoted in bold.

**Table S6. Epitope validation performance metrics.**

| | Precision | Recall | F1 |
|---|---|---|---|
| Pepsickle | 0.766 | **0.828** | **0.796** |
| NetChop | 0.671 | 0.747 | 0.707 |
| PCPS | 0.656 | 0.619 | 0.637 |
| PCleavage | **0.834** | 0.182 | 0.298 |

**Table S7. Command line computational performance.**

| | Model | |
|---|---|---|
| | pepsickle | NetChop 3.1 |
| epitope | 158m 21s | 542m 40s |
| in-vitro | 154m 46s | 260m 50s |

Time performances are based on processing time for the whole human proteome (see methods).

**Table S8. Amino acid feature matrix.**

| Amino Acids | One -Hot encoded ID's | | | | | | | | | | | | | | | | | | | | Physical Properties | | | |
|---|---|---|---|---|---|---|---|---|---|---|---|---|---|---|---|---|---|---|---|---|---|---|---|---|
| | A | C | D | E | F | G | H | I | K | L | M | N | P | Q | R | S | T | V | W | Y | Polarity (pI) | Molecular Volume | Hydrophobicity (cos -theta) | Confromational Entropy |
| A | 1 | 0 | 0 | 0 | 0 | 0 | 0 | 0 | 0 | 0 | 0 | 0 | 0 | 0 | 0 | 0 | 0 | 0 | 0 | 0 | 6 | 56.15265 | -0.495 | -2.4 |
| C | 0 | 1 | 0 | 0 | 0 | 0 | 0 | 0 | 0 | 0 | 0 | 0 | 0 | 0 | 0 | 0 | 0 | 0 | 0 | 0 | 5.07 | 69.61701 | 0.081 | -4.7 |
| D | 0 | 0 | 1 | 0 | 0 | 0 | 0 | 0 | 0 | 0 | 0 | 0 | 0 | 0 | 0 | 0 | 0 | 0 | 0 | 0 | 2.77 | 70.04515 | 9.573 | -4.5 |
| E | 0 | 0 | 0 | 1 | 0 | 0 | 0 | 0 | 0 | 0 | 0 | 0 | 0 | 0 | 0 | 0 | 0 | 0 | 0 | 0 | 3.22 | 86.35615 | 3.173 | -5.2 |
| F | 0 | 0 | 0 | 0 | 1 | 0 | 0 | 0 | 0 | 0 | 0 | 0 | 0 | 0 | 0 | 0 | 0 | 0 | 0 | 0 | 5.48 | 119.722 | -0.37 | -4.9 |
| G | 0 | 0 | 0 | 0 | 0 | 1 | 0 | 0 | 0 | 0 | 0 | 0 | 0 | 0 | 0 | 0 | 0 | 0 | 0 | 0 | 5.97 | 37.80307 | 0.386 | -1.9 |
| H | 0 | 0 | 0 | 0 | 0 | 0 | 1 | 0 | 0 | 0 | 0 | 0 | 0 | 0 | 0 | 0 | 0 | 0 | 0 | 0 | 7.59 | 97.94236 | 2.029 | -4.4 |
| I | 0 | 0 | 0 | 0 | 0 | 0 | 0 | 1 | 0 | 0 | 0 | 0 | 0 | 0 | 0 | 0 | 0 | 0 | 0 | 0 | 6.02 | 103.6644 | -0.528 | -6.6 |
| K | 0 | 0 | 0 | 0 | 0 | 0 | 0 | 0 | 1 | 0 | 0 | 0 | 0 | 0 | 0 | 0 | 0 | 0 | 0 | 0 | 9.74 | 102.7783 | 2.101 | -7.5 |
| L | 0 | 0 | 0 | 0 | 0 | 0 | 0 | 0 | 0 | 1 | 0 | 0 | 0 | 0 | 0 | 0 | 0 | 0 | 0 | 0 | 5.98 | 102.7545 | -0.342 | -6.3 |
| M | 0 | 0 | 0 | 0 | 0 | 0 | 0 | 0 | 0 | 0 | 1 | 0 | 0 | 0 | 0 | 0 | 0 | 0 | 0 | 0 | 5.74 | 103.928 | -0.324 | -6.1 |
| N | 0 | 0 | 0 | 0 | 0 | 0 | 0 | 0 | 0 | 0 | 0 | 1 | 0 | 0 | 0 | 0 | 0 | 0 | 0 | 0 | 5.41 | 76.56687 | 2.354 | -4.7 |
| P | 0 | 0 | 0 | 0 | 0 | 0 | 0 | 0 | 0 | 0 | 0 | 0 | 1 | 0 | 0 | 0 | 0 | 0 | 0 | 0 | 6.3 | 71.24858 | -0.322 | -0.8 |
| Q | 0 | 0 | 0 | 0 | 0 | 0 | 0 | 0 | 0 | 0 | 0 | 0 | 0 | 1 | 0 | 0 | 0 | 0 | 0 | 0 | 5.65 | 88.62562 | 2.176 | -5.5 |
| R | 0 | 0 | 0 | 0 | 0 | 0 | 0 | 0 | 0 | 0 | 0 | 0 | 0 | 0 | 1 | 0 | 0 | 0 | 0 | 0 | 10.76 | 110.5867 | 4.383 | -6.9 |
| S | 0 | 0 | 0 | 0 | 0 | 0 | 0 | 0 | 0 | 0 | 0 | 0 | 0 | 0 | 0 | 1 | 0 | 0 | 0 | 0 | 5.68 | 55.89516 | 0.936 | -4.6 |
| T | 0 | 0 | 0 | 0 | 0 | 0 | 0 | 0 | 0 | 0 | 0 | 0 | 0 | 0 | 0 | 0 | 1 | 0 | 0 | 0 | 5.6 | 72.0909 | 0.853 | -5.1 |
| V | 0 | 0 | 0 | 0 | 0 | 0 | 0 | 0 | 0 | 0 | 0 | 0 | 0 | 0 | 0 | 0 | 0 | 1 | 0 | 0 | 5.96 | 86.28358 | -0.308 | -4.6 |
| W | 0 | 0 | 0 | 0 | 0 | 0 | 0 | 0 | 0 | 0 | 0 | 0 | 0 | 0 | 0 | 0 | 0 | 0 | 1 | 0 | 5.89 | 137.5186 | -0.27 | -4.8 |
| Y | 0 | 0 | 0 | 0 | 0 | 0 | 0 | 0 | 0 | 0 | 0 | 0 | 0 | 0 | 0 | 0 | 0 | 0 | 0 | 1 | 5.66 | 121.5862 | 1.677 | -5.4 |
| * | 0 | 0 | 0 | 0 | 0 | 0 | 0 | 0 | 0 | 0 | 0 | 0 | 0 | 0 | 0 | 0 | 0 | 0 | 0 | 0 | 7.5 | 0 | 1.689157 | 0 |
| B | 0 | 0 | 1 | 0 | 0 | 0 | 0 | 0 | 0 | 0 | 0 | 1 | 0 | 0 | 0 | 0 | 0 | 0 | 0 | 0 | 4.09 | 73.30601 | 5.964 | -4.6 |
| Z | 0 | 0 | 0 | 1 | 0 | 0 | 0 | 0 | 0 | 0 | 0 | 0 | 0 | 1 | 0 | 0 | 0 | 0 | 0 | 0 | 4.44 | 87.49089 | 2.675 | -5.35 |
| J | 0 | 0 | 0 | 0 | 0 | 0 | 0 | 1 | 0 | 1 | 0 | 0 | 0 | 0 | 0 | 0 | 0 | 0 | 0 | 0 | 6 | 103.2094 | -0.426 | -6.45 |
| U | 0 | 1 | 0 | 0 | 0 | 0 | 0 | 0 | 0 | 0 | 0 | 0 | 0 | 0 | 0 | 0 | 0 | 0 | 0 | 0 | 5.07 | 69.61701 | 0.081 | -4.7 |
| X | 1 | 1 | 1 | 1 | 1 | 1 | 1 | 1 | 1 | 1 | 1 | 1 | 1 | 1 | 1 | 1 | 1 | 1 | 1 | 1 | 6.008095 | 88.55829 | 0.6195 | -4.845 |

Encoded feature matrix including bit vector notation and physical/chemical properties for each standard amino acid and recognized ambiguous amino acids.

**Table S9. In vitro model performance on epitope validation data.**

| Proteasome mode | Sensitivity | Specificity | AUC |
|---|---|---|---|
| Constitutive | 69.85% | 51.49% | 0.650 |
| Immuno | 54.54% | 73.71% | 0.679 |
