## Supplementary_Figures for "Pepsickle rapidly and accurately predicts proteasomal cleavage sites for improved neoantigen identification"

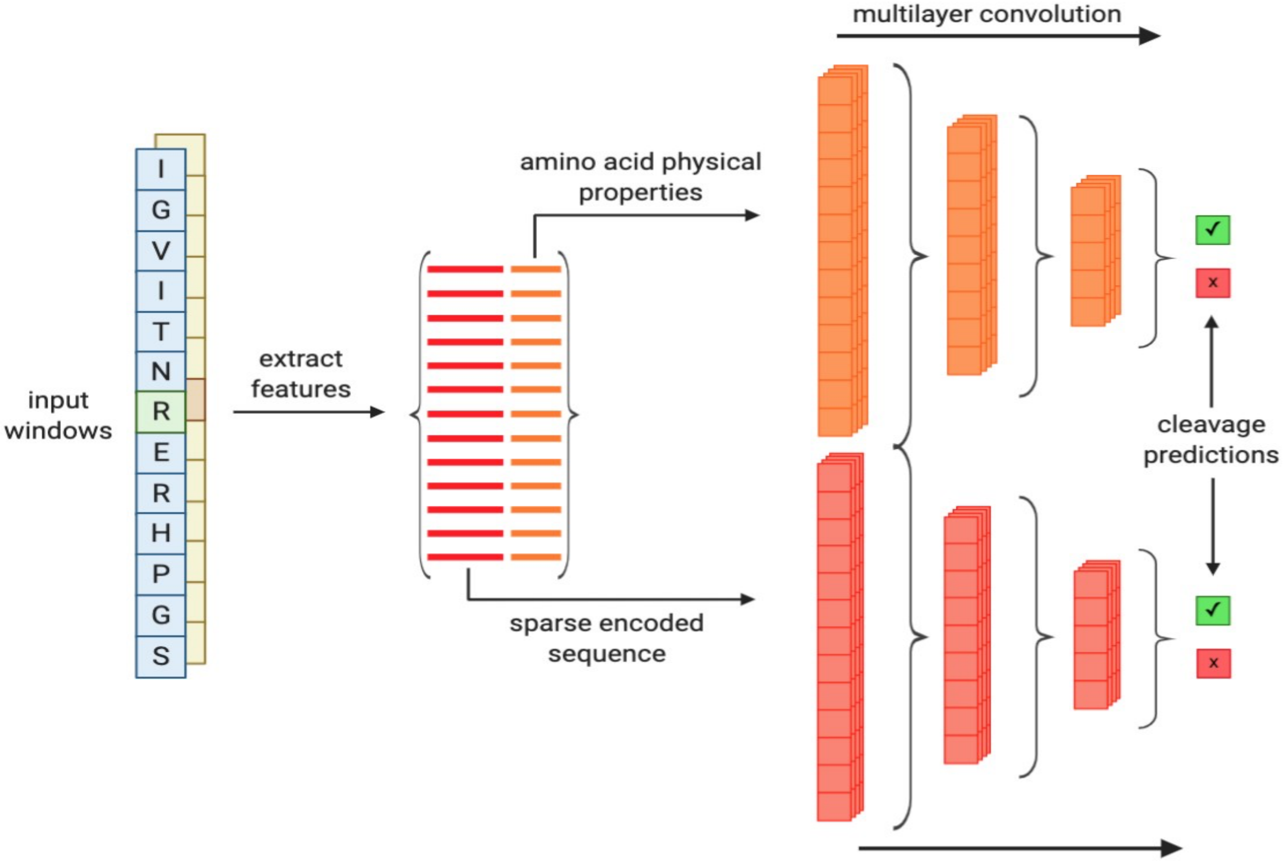


**Figure S1. Epitope consensus model layout.** Features from amino acid windows in the epitope datasets were extracted to identify the one-hot encoded amino acid sequences, as well as the physical properties at each window position. One-hot encoded sequences were fed directly into the first layer of the deep learning model, while physical properties underwent a 1D convolution (span = 3) across each property prior to first layer input. For each internal layer, ReLU activation functions were used with 20% dropout. For final layers, log(SoftMax) was used to give class probability outputs. For exact layer numbers and sizes based on input window size see **Table S2 & Table S3.**

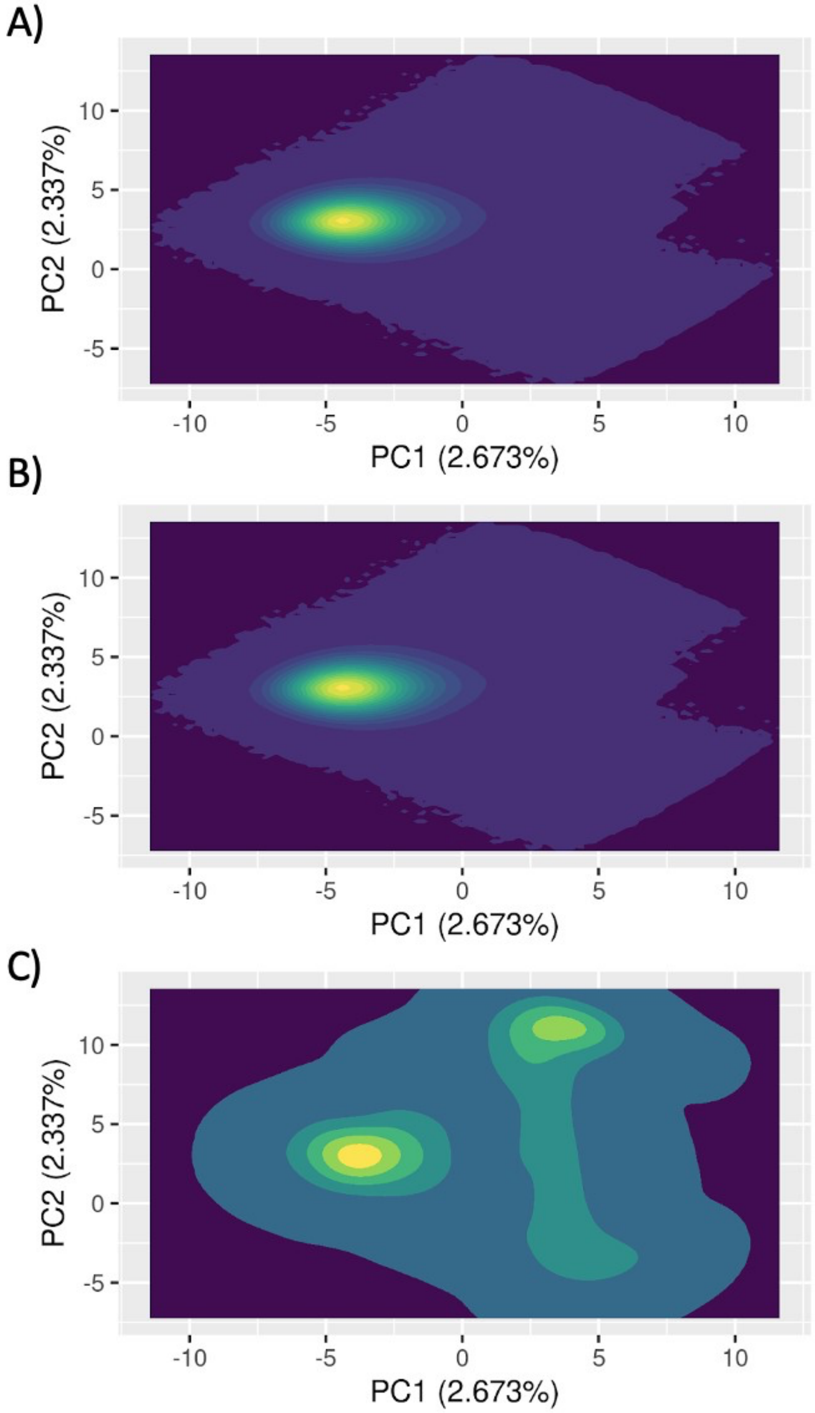


**Figure S2. Training set sample densities compared to human background. A)** Principle components were constructed using the physical property values across 21 amino acid windows generated from all proteins in the human proteome. Using the first and second principle components (PC1 and PC2, respectively), sample density was calculated and plotted in PCA space. **B)** The density distribution for all 21 amino acid windows in the epitope based training set are shown using the same encoding and PCA approach. **C)** The density distribution for all *in vitro* based training examples are shown based on the same encoding approach.

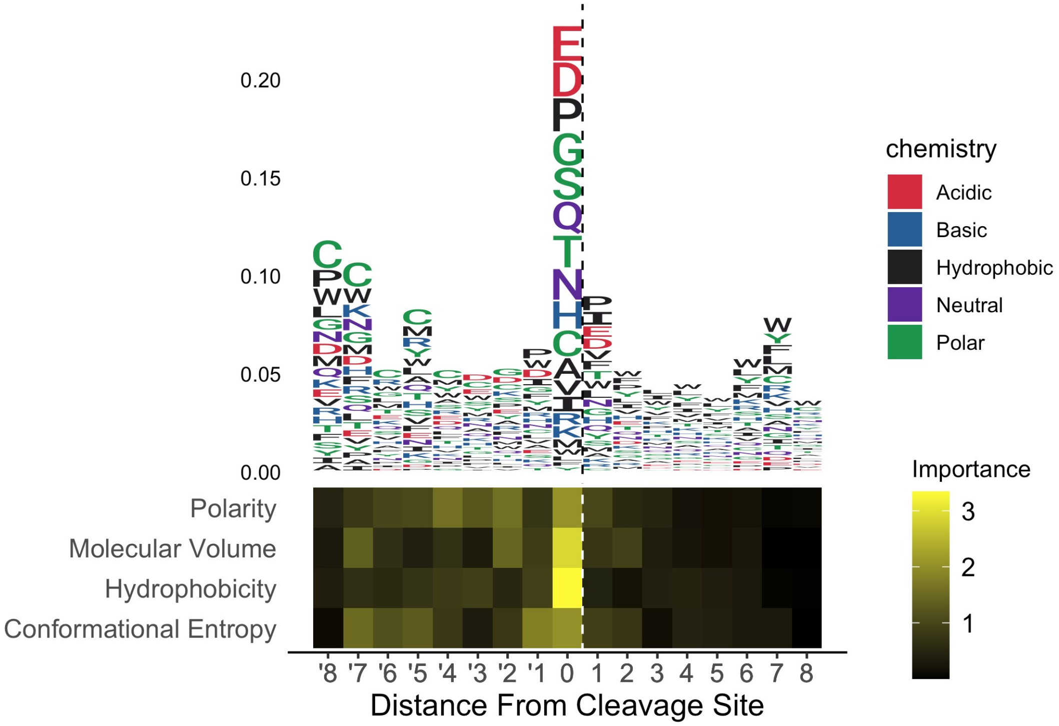


**Figure S3. Epitope consensus model feature importances.** Feature importances were calculated as the absolute values of the model saliencies for the sequence identities (top) and chemical properties (bottom) at each given position in the input window of our 17 amino acid consensus model. For sequences, the total height of each bar corresponds to overall importance of a given position in the model, while the height of each letter corresponds to importance of the corresponding amino acid at that position. Chemical property feature importance is indicated by color gradient from most important (yellow) to least important (black).

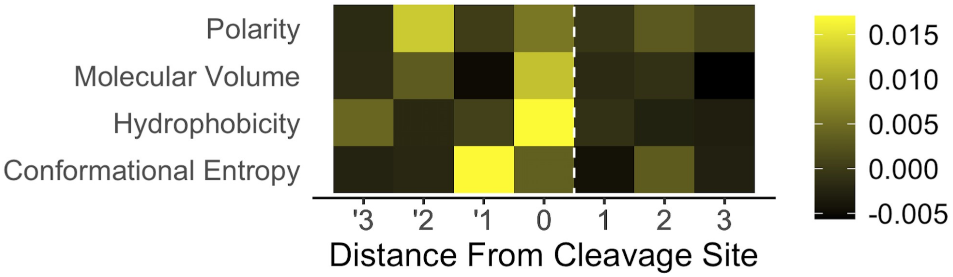


**Figure S4. Chemical property feature importances for *in vitro* digestion model.** Feature importances were calculated as the normalized absolute values of the model weights for chemical properties at each given position in the input window of our 7 amino acid digestion based *in vitro* model. Feature importance is indicated by color gradient from most important (yellow) to least important (black).

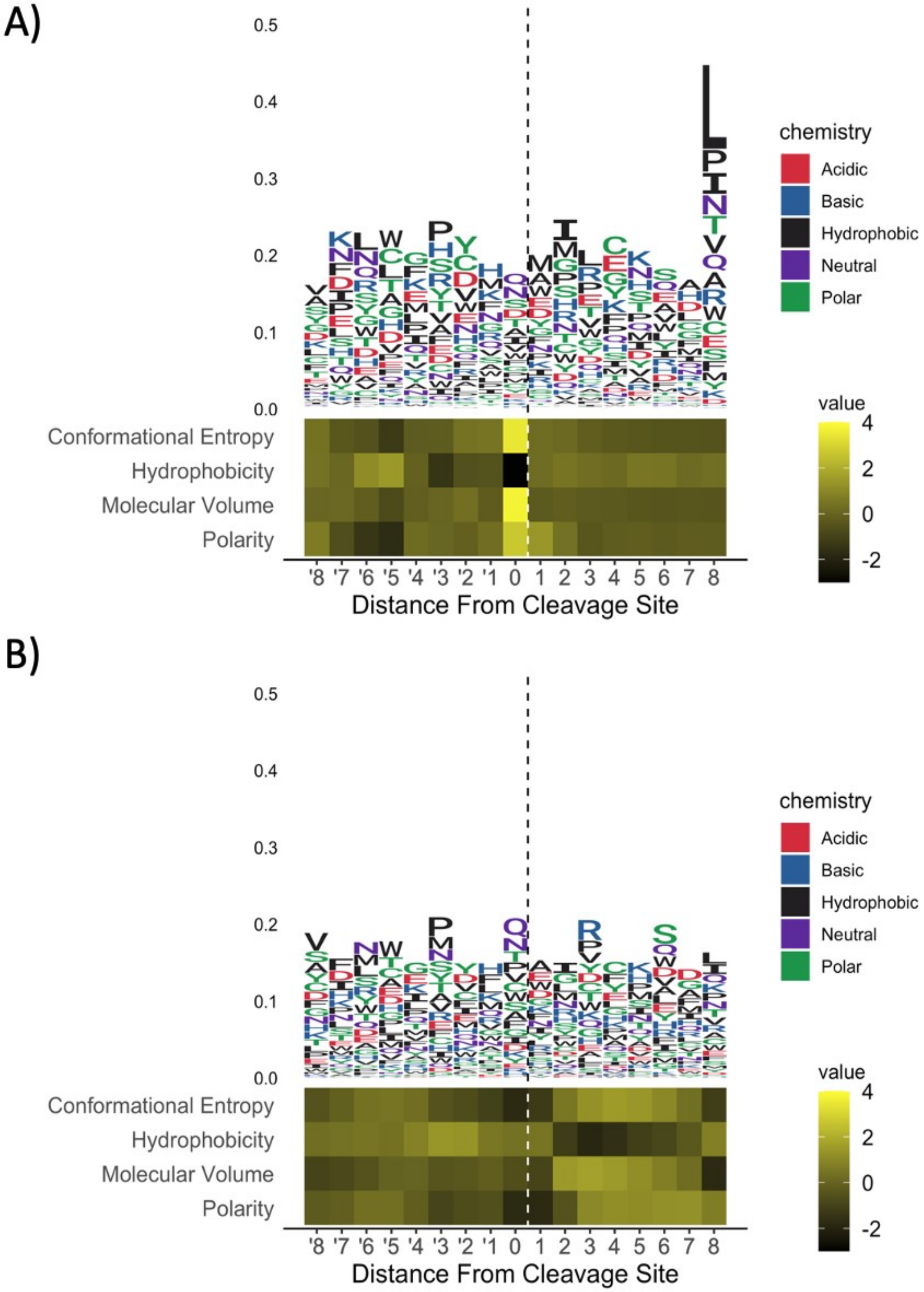


**Figure S5. Epitope training set features. A)** Amino acid identities (top) and chemical properties (bottom) from positive cleavage windows were plotted as the average frequency (sequence) or average normalized value (chemical properties) across all amino acids at a given position. **B)** Non-cleavage windows were plotted using the same schema and ranges used for cleavage events.

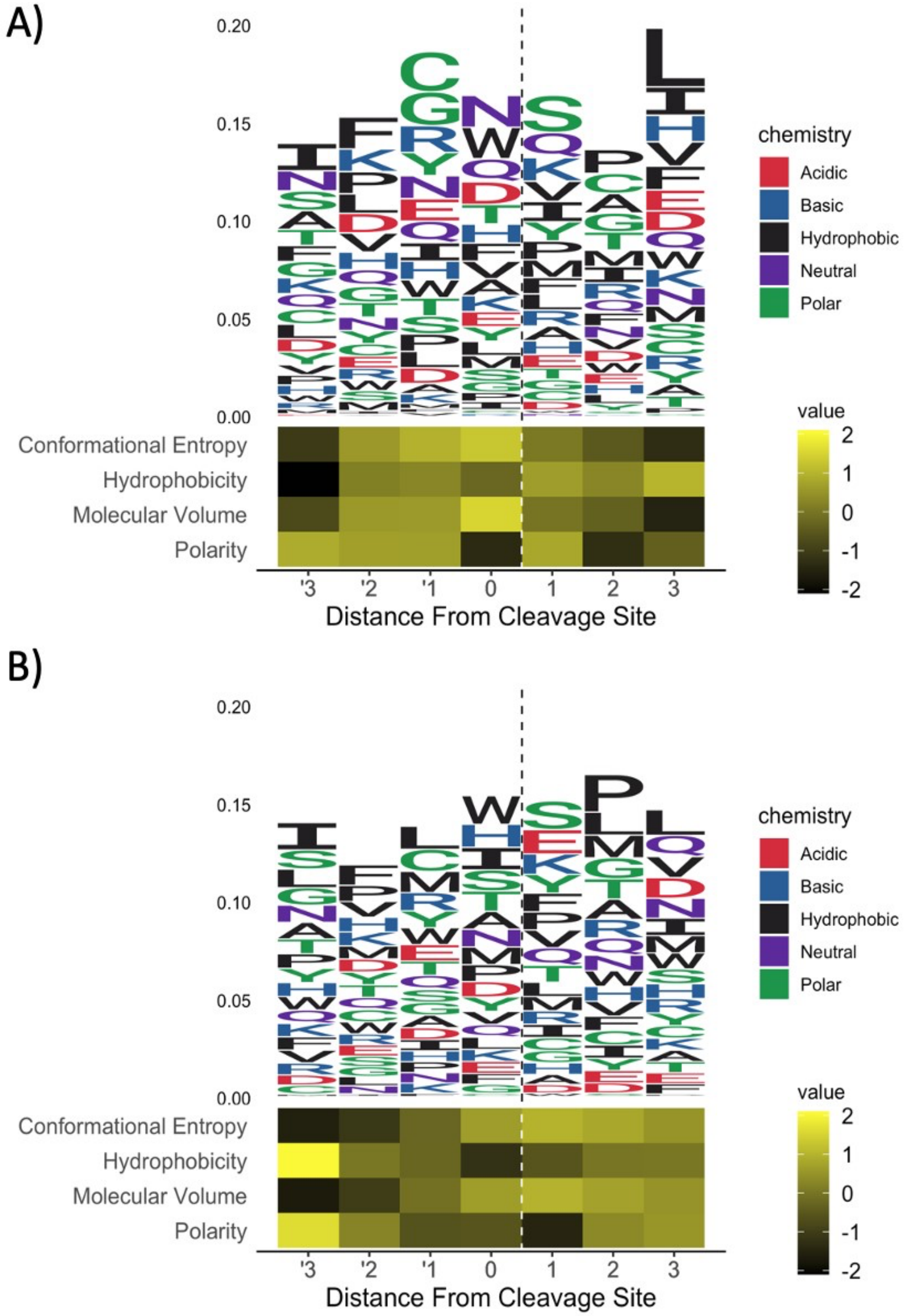


**Figure S6. Digestion training set features. A)** Amino acid identities (top) and chemical properties (bottom) from positive cleavage windows were plotted as the average frequency (sequence) or average normalized value (chemical properties) across all amino acids at a given position. **B)** Non-cleavage windows were plotted using the same schema and ranges used for cleavage events.
